# An EPHB4-RASA1 signaling complex inhibits shear stress-induced Ras-MAPK activation in lymphatic endothelial cells to promote the development of lymphatic vessel valves

**DOI:** 10.1101/2023.11.22.568378

**Authors:** Di Chen, David Wiggins, Eva M. Sevick, Michael J. Davis, Philip D. King

## Abstract

EPHB4 is a receptor protein tyrosine kinase that is required for the development of lymphatic vessel (LV) valves. We show here that EPHB4 is necessary for the specification of LV valves, their continued development after specification, and the maintenance of LV valves in adult mice. EPHB4 promotes LV valve development by inhibiting the activation of the Ras-MAPK pathway in LV endothelial cells (LEC). For LV specification, this role for EPHB4 depends on its ability to interact physically with the p120 Ras-GTPase-activating protein (RASA1) that acts as a negative regulator of Ras. Through physical interaction, EPHB4 and RASA1 dampen oscillatory shear stress (OSS)-induced Ras-MAPK activation in LEC, which is required for LV specification. We identify the Piezo1 OSS sensor as a focus of EPHB4-RASA1 regulation of OSS-induced Ras-MAPK signaling mediated through physical interaction. These findings contribute to an understanding of the mechanism by which EPHB4, RASA1 and Ras regulate lymphatic valvulogenesis.

## INTRODUCTION

Ephrin receptors (EPHR) constitute a large family of receptor protein tyrosine kinases (RTK) in mammals ^1–3^. There are nine EPHR class A receptors (EPHA) that recognize five different glycophosphatidylinositol-linked cell surface Ephrin A ligands, and five EPHR class B receptors (EPHB) that recognize three different transmembrane Ephrin B ligands^4,5^. The extracellular region of EPHR comprises of a ligand-binding domain, a cysteine-rich region and two fibronectin domains. The intracellular domain of EPHR contains a juxtamembrane alpha-helical region with regulatory tyrosine residues followed by a protein tyrosine kinase domain and a terminal sterile alpha motif domain^1–5^. EPHR regulate diverse cellular processes such as cell migration, survival, proliferation and differentiation in multiple cell types and tissues^1–3^. In the cardiovascular system, EPHB4 in particular has emerged as essential for both its normal development and function^6^.

Genetically engineered mice that lack EPHB4 constitutively die in mid-gestation as a consequence of impaired developmental angiogenesis, wherein primitive vascular plexuses formed through the process of vasculogenesis are not remodeled properly into hierarchical arterial-capillary-venous networks^7,8^. In addition, with the use of conditional EPHB4-deficient mice, it has been demonstrated that EPHB4 is required for the development of intraluminal valves in collecting lymphatic vessels that are necessary for propulsive lymph flow^9^, and for the formation of lymphovenous valves that prevent backflow of venous blood into the lymphatic vascular system^10^. Furthermore, EPHB4 has been shown to be necessary for retinal blood vascular angiogenesis in newborns^11^, for pathological angiogenesis toward solid tumors in adults^11^, and for the normal functioning of the cardiac capillary network in adults^12^. Further evidence of the important function of EPHB4 in the vasculature has come from human genetic studies. Germline inactivating mutations of *EPHB4* in man are responsible for one form of an autosomal dominant vascular disease named capillary malformation-arteriovenous malformation (CM-AVM), the pathognomonic feature of which is the presence of one or more cutaneous CM, with additional AVM in one-third of patients^13^. Germline inactivating mutations of *EPHB4* have also been identified as causative of Vein of Galen Arteriovenous malformation (VGAM)^14,15^ and for lymphatic vessel anomalies including lymphatic related hydrops fetalis (LRHF) that involves lymph accumulation in the fetus^10,16^, lymphedema in which lymph accumulates in peripheral tissues^16,17^, and central conducting lymphatic anomaly^18^.

Until recently, the signaling mechanisms downstream of EPHB4 that permit normal cardiovascular development and function in vivo had not been studied. Studies performed in cell lines in vitro demonstrated that EPHB4 is able to activate the Ras-mitogen-activated protein kinase (MAPK) pathway^19^. However, the ability of EPHB4 to activate this pathway is cell context dependent. Whereas in neuronal cells and transformed cells, EPHB4 can trigger Ras-MAPK signaling, in endothelial cells (EC), EPHB4 functions to inhibit activation of this pathway induced by the engagement of other RTK, including vascular endothelial growth factor receptor-2 (VEGFR2) and Tie-2^19,20^. The ability of EPHB4 to dampen Ras-MAPK activation in EC in vitro is dependent upon the expression of the p120 Ras GTPase-activating protein (p120 RasGAP), also known as Ras-activating protein 1 (RASA1)^19^. RASA1 is one member of a family of ten RasGAPs that negatively regulate the Ras-MAPK pathway by increasing the ability of Ras to hydrolyze bound GTP to GDP, thereby resulting in Ras inactivation^21^.

In mice, *Rasa1* loss of function mutations essentially phenocopy *Ephb4* loss of function mutations. Thus, embryos constitutively deficient in functional RASA1 die in mid-gestation from impaired developmental angiogenesis^22,23^, whereas later induced disruption of *Rasa1* in the vasculature results in failed development of lymphatic valves^24^, lymphovenous valves^25^, and venous valves^25^. After birth, RASA1 is also necessary for retinal angiogenesis in newborns and tumor angiogenesis in adults^26^, and for the maintenance of lymphatic valves^24,27^ and venous valves^25^. In humans, loss of function mutations in *RASA1* are responsible for all remaining cases of CM-AVM not caused by *EPHB4* mutations^28,29^. In addition, loss of function mutations of *RASA1* are causative of some cases of VGAM^28–30^ and Parkes-Weber syndrome that involves diffuse CM and AVM^28,29,31^, and lymphatic dysfunction including chylous ascites^29^ and chylothorax^32^ in which lymph fluid accumulates in the peritoneal and pleural cavities respectively, lymphatic malformation^31,33^, LRHF^34^, and lymphedema^35^. These findings in mouse and man are consistent with the notion that EPHB4 and RASA1 perform the same function in the same signaling pathway to regulate vascular development and function^6^.

Mechanistically, loss of EPHB4 or RASA1 during developmental angiogenesis results in dysregulated Ras-MAPK signaling in EC that drives increased expression and abundance of collagen IV proline and lysine hydroxylases in the endoplasmic reticulum (ER)^11,26^. This is thought to lead to over-hydroxylation of collagen IV monomers in the EC ER that impacts their folding and assembly into trimeric protomers, in which form collagen IV is normally secreted from EC and deposited in basement membranes. Consequently, EC undergo apoptotic death due to an unresolved unfolded protein response (UPR) and/or an inability to attach properly to an incompletely formed basement membrane^11,26^. Consistent with this model, drugs that inhibit the MAPK pathway or inhibit proline and lysine hydroxylases, and drugs that promote folding of collagen IV in the EC ER, each rescue EC collagen IV accumulation in the ER, EC apoptosis and developmental angiogenesis in induced EPHB4-deficient or RASA1-deficient embryos^11,26^. These findings substantiate the notion that EPHB4 and RASA1 communicate with one another to negatively regulate Ras-MAPK signaling in EC necessary for normal vascular development in vivo^6^.

RASA1 is known to interact physically with EPHB4 through RASA1 SH2 domain-mediated recognition of two tyrosine residues in the EPHB4 JM region that are phosphorylated during the course of EPHB4 signaling^11,36^. Therefore, a simple a priori model to account for EPHB4-RASA1 communication in the regulation of Ras-MAPK signaling during angiogenesis is that EPHB4 serves to recruit RASA1 to the plasma membrane, allowing its juxtaposition to Ras-GTP necessary for Ras inactivation. To address this, we recently generated an EPHB4 knockin mouse that expresses a quadruple point mutant form of EPHB4 (Y590F/P593G/Y596F/P599G) named EPHB4 2YP, that is specifically unable to bind RASA1 yet retains PTK catalytic activity^11^. Surprisingly, developmental, neonatal, and pathological blood vascular angiogenesis, and developmental lymphangiogenesis were all intact in these mice^11^. This finding indicates that physical interaction between EPHB4 and RASA1 is not required for these events.

Although EPHB4 is known to be required for lymphatic valve development^9,37^, at which stage in lymphatic valvulogenesis EPHB4 plays a role and whether EPHB4 is necessary for valve maintenance in adults is unknown. Similarly, whether the role of EPHB4 in valvulogenesis relates to an ability to regulate Ras-MAPK signaling through physical interaction with RASA1 is unknown. In this study, we show that EPHB4 is required for lymphatic valve specification, continued valve development after specification, and valve maintenance in adults. In addition, we show that in the former two cases, EPHB4 functions by limiting the activation of the Ras-MAPK pathway in lymphatic EC (LEC). Most importantly, we further show that physical interaction between EPHB4 and RASA1 is necessary to limit Ras-MAPK activation induced by the Piezo1 shear stress sensor in LEC to allow lymphatic valve specification.

## RESULTS

### EPHB4 is required for the specification and continued development of LV valves

It has been shown previously that genetic disruption of *Ephb4* in EC using a *pdgfb-ert2cre* driver results in impaired postnatal development of LV valves^9^. To confirm these findings and demonstrate that impaired valve development results from the loss of EPHB4 in LEC, we generated *Ephb4^fl/fl^* mice with and without the LEC-specific *Prox1-ert2cre* driver^38^. Pups were administered tamoxifen (TM) on day 4 after birth (P4) and valve number in collecting LV was later assessed by whole mount staining for PROX1 that is expressed at higher levels in lymphatic valve LEC than lumenal wall LEC ^39,40^ (Figure 1A). One week after TM administration, there were much fewer valves in mesenteric and diaphragmatic collecting LV of *Cre−*positive pups than *Cre*-negative littermate controls (Figure 1B, C). Thus, expression of EPHB4 within LEC is necessary for the postnatal development of LV valves.

**Figure 1.**
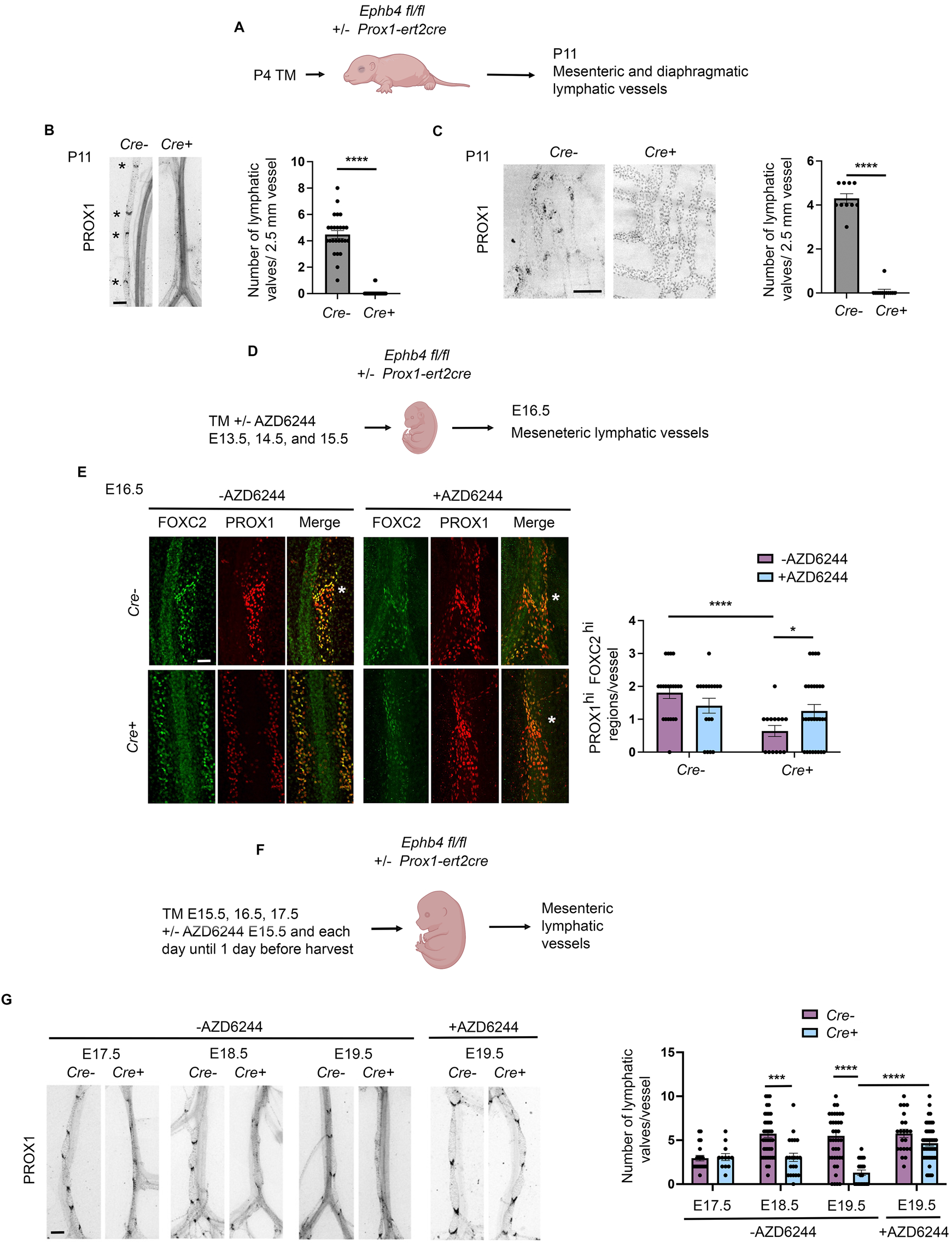
LV valve specification and development in induced LEC-specific EPHB4-deficient mice. (A) Schematic showing design of experiments in (B and C). (B) At left are shown representative images of PROX1 staining of mesenteric LV. Scale bar=200 µm. PROX1^hi^ valves are indicated with asterisks. At right is shown the mean +/− 1 SEM of valve number in mesenteric LV (*Cre−*, n=25 determinations from 5 mice; *Cre+*, n=44 determinations from 9 mice). **** *P*<0.0001, two-tailed Mann-Whitney test. (C) At left are shown representative images of PROX1 staining of diaphragmatic LV. Scale bar=200 µm. At right is shown the mean +/− 1 SEM of valve number in diaphragmatic LV (*Cre−*, n=10 determinations from 2 mice; *Cre+*, n=12 determinations from 2 mice). **** *P*<0.0001, two-tailed Mann-Whitney test. (D) Schematic showing design of experiments in (E). (E) At left are shown representative images of PROX1 and FOXC2 staining of mesenteric LV. Scale bar=100 µm. Asterisks in merged images indicate sites of high FOXC2 and PROX1 staining. At right is shown the mean +/− 1 SEM of PROX1^hi^ FOXC2^hi^ regions per LV without AZD6244 treatment (*Cre−*, n=21 vessels from 3 embryos; *Cre+*, n=14 vessels from 3 embryos) and with AZD6244 treatment (*Cre−*, n=17 vessels from 2 embryos; *Cre+*, n=28 vessels from 3 embryos). * *P*<0.05, **** *P*<0.0001, two-tailed Student’s 2-sample t-test. (F) Schematic showing design of experiments in (G). (G) Representative images of PROX1 staining are shown at left. Scale bar=200 µm. At right is shown the mean +/− 1 SEM of numbers of valves per LV without AZD6244 treatment at E17.5 (*Cre−*, n=20 vessels from 5 embryos; *Cre+*, n=12 vessels from 2 embryos), E18.5 (*Cre−*, n=42 vessels from 6 embryos; *Cre+*, n=21 vessels from 3 embryos) and E19.5 (*Cre−*, n=41 vessels from 8 embryos; *Cre+*, n=20 vessels from 3 embryos), and with AZD6244 treatment at E19.5 (*Cre−*, n=20 vessels from 3 embryos; *Cre+*, n=46 vessels from 8 embryos). *** *P*<0.001, **** *P*<0.0001, two-tailed Mann-Whitney test.

At which stage of LV valve development EPHB4 functions is unknown. EPHB4 could be necessary for initial valve specification, continued valve development after specification, or both. To address this, we examined valve development in embryonic mesenteric LV, where the times and stages of valve development are well defined ^39,40^. In embryonic mesenteric LV, valve specification initiates at E15 and is marked by increased expression of the FOXC2 and PROX1 transcription factors ^39,40^. Therefore, to determine if EPHB4 is necessary for LV specification, we administered TM to E13.5 *Ephb4^fl/fl^ Prox1-ert2cre* embryos and examined LV valve specification by whole mount staining of mesenteric LV at E16.5 using FOXC2 and PROX1 antibodies (Figure 1D). At E16.5, the frequency of PROX1^hi^FOXC2^hi^ LV valve-forming regions in mesenteric LV was significantly lower in *Ephb4^fl/fl^ Prox1-ert2cre* embryos compared to littermate *Ephb4^fl/fl^* embryos (Figure 1E). Therefore, EPHB4 promotes LV valve specification.

To examine if EPHB4 is necessary for continued LV valve development after initial specification, we delayed administration of TM to *Ephb4^fl/fl^ Prox1-ert2cre* embryos until E15.5 (Figure 1F). Analysis of valve number in mesenteric LV at E17.5 did not reveal significant differences between *Ephb4^fl/fl^ Prox1-ert2cre* and *Ephb4^fl/fl^* littermate control embryos (Figure 1G). However, at E18.5 and E19.5, *Ephb4^fl/fl^ Prox1-ert2cre* embryos had significantly reduced numbers of valves compared to controls (Figure 1G). These findings are in support of a continued role for EPHB4 in valve development after initial specification.

In EC specifically, EPHB4 is thought to limit activation of the Ras-MAPK pathway through functional interaction with RASA1^6,11,19,20^. Therefore, we asked if impaired LV valve specification and continued valve development in LEC-specific EPHB4-deficient embryos could be rescued by inhibition of the Ras-MAPK pathway. For this purpose, we administered the MAPK pathway inhibitor, AZD6244, to embryos at the same time as TM. In experiments that examined valve specification, AZD6244 could rescue increased expression of FOXC2 and PROX1 in *Ephb4^fl/fl^ Prox1-ert2cre* embryos as assessed at E16.5 (Figure 1E). Likewise, AZD6244 was able to rescue continued valve development as assessed at E19.5 (Figure 1G). These findings are consistent with the notion that in the absence of EPHB4, dysregulated activation of the Ras MAPK pathway in LEC is responsible for both impaired valve specification and impaired continued valve development post-specification.

### EPHB4 is required for LV valve maintenance and function in adult mice

Whether EPHB4 is necessary for maintaining LV valves in adults has not been addressed. To examine this, we administered TM to adult *Ephb4^fl/fl^ Prox1-ert2cre* mice. One week later, we examined the extent of activation of the Ras-MAPK pathway by immunostaining of sections of popliteal collecting LV using an anti-phospho-ERK MAPK antibody (Figure 2A). At this time, the percentage of LV valve LEC with detectable pERK signals was consistently higher in *Ephb4^fl/fl^ Prox1-ert2cre* mice compared to littermate *Ephb4^fl/fl^* mice (Figure 2B). This increase was apparent in both LV valve leaflet LEC and in LEC of the valve sinus wall and is consistent with the role of EPHB4 as a negative regulator of the Ras-MAPK pathway in LEC.

**Figure 2.**
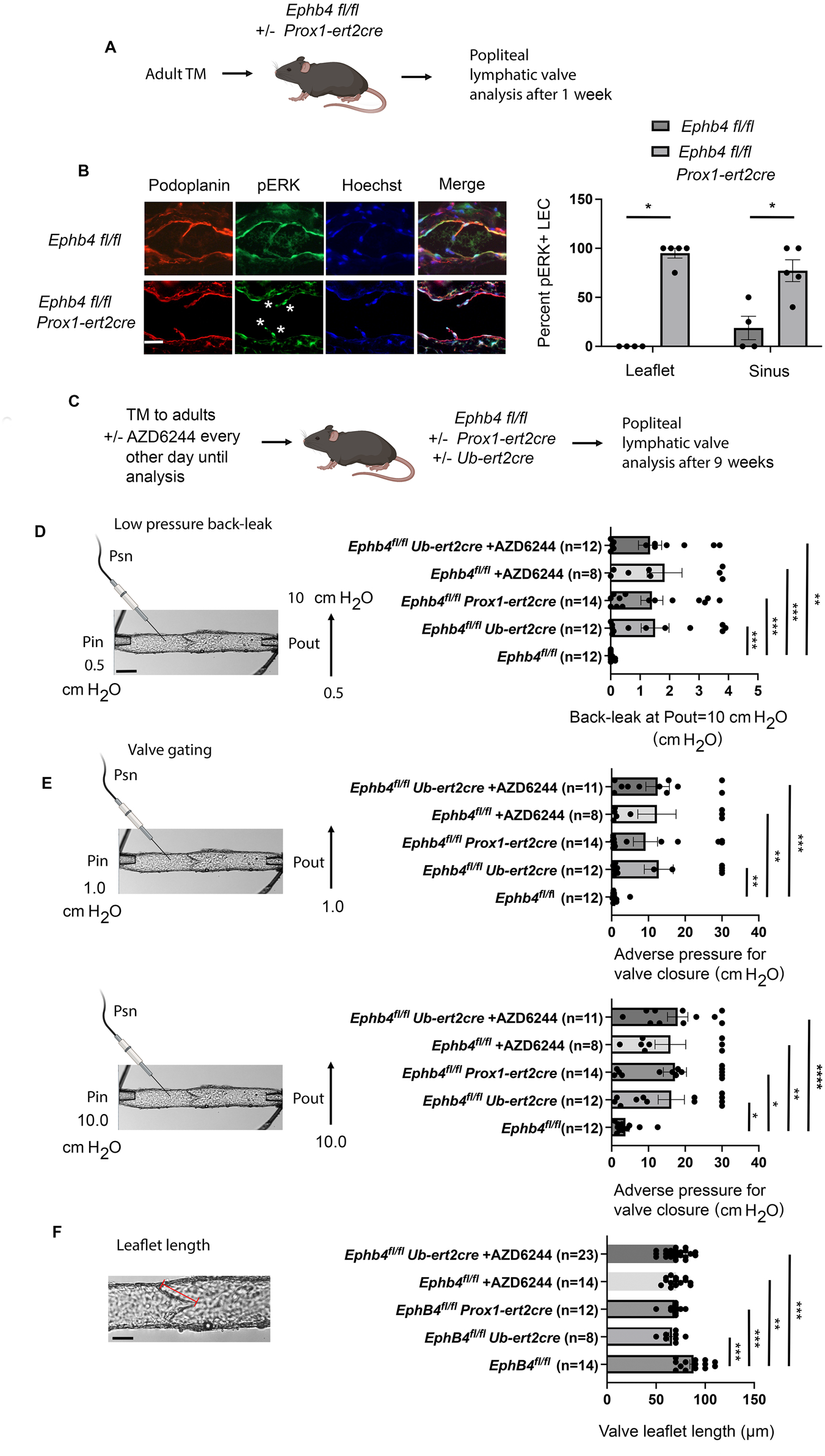
LV valve function in induced LEC-specific adult EPHB4-deficient mice. (A) Schematic showing design of experiments in (B). (B) At left are shown representative images of sections of popliteal LV valve regions stained with the indicated antibodies or Hoecsht. Scale bar=50 µm. Note pERK+ LEC in valve leaflets of *Cre*+ mouse, marked with asterisks. At right is shown the mean +/− 1 SEM of the percent pERK+ LEC in LV valve leaflets and sinuses (*Cre−*, n=4 valve regions from 3 mice; *Cre+*, n=5 valve regions from 4 mice). * *P*<0.05, two-tailed Mann-Whitney test. (C) Schematic showing design of experiments in (D-F). (D) Image at left shows design of low pressure back-leak tests. Scale bar=100 µm. Graph shows mean +/− 1 SEM of the amount of pressure back-leak at a Pout of 10 cm H_2_O. Numbers of tested valves are indicated in parentheses and were obtained from the following numbers of mice: *Ephb4 fl/fl*, n=5; *Ephb4 fl/fl Ub-ert2cre*, n=4; *Ephb4 fl/fl Prox1-ert2cre*, n=5; *Ephb4 fl/fl +*AZD6244 n=3; *Ephb4 fl/fl Ub-ert2cre* + AZD6244, n=4). ** *P*<0.01, *** *P*<0.001, two-tailed Mann-Whitney test. (E) Images at left show design of valve gating tests. Graphs shows mean +/− 1 SEM of the amount of adverse pressure required to close valves at a Pin of 1 or 10 cm H_2_O. Numbers of tested valves are indicated in parentheses and were obtained from the same number of mice as indicated in (D). * *P*<0.05, ** *P*<0.01, *** *P*<0.001, **** *P*<0.0001, two-tailed Mann-Whitney test. (F) The red bar in the valve image indicates the distance measured in the determination of valve leaflet length. Scale bar=50 µm. The graph shows the mean +/− 1 SEM of valve leaflet lengths. Numbers of valve leaflets measured are indicated in parentheses and were obtained from the same number of mice as indicated in (D), except for *Ephb4 fl/fl* controls without AZD6244 treatment that include *Ephb4 fl/fl* controls not treated with TM in Figure 4E (n=9 mice total). ** *P*<0.01, *** *P*<0.001, two-tailed Mann-Whitney test.

To assess the consequence of EPHB4 loss upon LV valve function in adults, we performed different valve function tests 9 weeks after TM administration (Figure 2C). Popliteal collecting LV were dissected from mice, were trimmed to contain a single valve, and were cannulated at both ends to permit manipulation of pressure upstream (Pin) and downstream (Pout) of the valve. In low-pressure back-leak tests, Pin was kept constant at 0.5 cm of H_2_O and Pout was elevated from 0.5 to 10 cm H_2_O. Throughout the Pout pressure elevation, any pressure back-leak across the valve was monitored using a Psn pipette inserted into the vessel lumen upstream of the valve (Figure 2D). As expected, valves from control *Ephb4^fl/fl^* mice prevented pressure back-leak up to Pout pressures of 10 cm of H_2_O. In contrast, most valves from *Ephb4^fl/fl^ Prox1-ert2cre* mice showed significant back-leak (Figure 2D). Similarly, the majority of valves from *Ephb4^fl/fl^* mice with a ubiquitously active *Ub-ert2cre* driver that were treated with TM 9 weeks previously showed significant back-leak (Figure 2D). In valve gating tests, Pin was held at 1 or 10 cm of H_2_O (which results in different vessel diameters at the start of the test), Pout was elevated, and the amount of adverse pressure required to close valves was determined (Figure 2E). In these assays, valves from control *Ephb4^fl/fl^* mice again behaved as expected. Most valves required relatively modest amounts of adverse pressure to close at either Pin pressure (<4 cmH_2_O). In contrast, valves from *Ephb4^fl/fl^ Prox1-ert2cre* and *Ephb4^fl/fl^ Ub-ert2cre* mice, on average, required increased amounts of adverse pressure for valve closure at both Pin pressures with a substantial percentage of valves unable to close at the highest pressure differential (30 cmH_2_O) that could be applied (Figure 2E). Therefore, EPHB4 is required to maintain LV valve function in adults. Determination of LV valve leaflet length in valve function assays revealed shorter leaflets in *Ephb4^fl/fl^ Prox1-ert2cre* and *Ephb4^fl/fl^ Ub-ert2cre* vessels compared to *Ephb4^fl/fl^* control vessels, which likely accounts for the observed valve dysfunction (Figure 2F). As determined by immunostaining and confocal imaging of valves, the shortened leaflet length was associated with reduced numbers of PROX1^hi^ LEC in leaflets (Figure S1).

To determine if LV valve dysfunction in induced EPHB4-deficient adult mice was a consequence of dysregulated Ras-MAPK signaling, we administered AZD6244 to *Ephb4^fl/fl^ Ub-ert2cre* mice and control *Ephb4^fl/fl^* control mice at the same time as TM and for every 2 days thereafter until valve functional analysis at 9 weeks. Administration of AZD6244 to *Ephb4^fl/fl^ Ub-ert2cre* mice did not significantly rescue valve function in either low-pressure back-leak or valve gating assays (Figure 2D, E). However, AZD6244 induced significant valve dysfunction in control *Ephb4^fl/fl^* mice in both assay types associated with leaflet shortening, which precludes straightforward interpretation of results (Figure 2D, E). This finding also indicates that in contrast to valve development, activation of the MAPK pathway is necessary to maintain LV valve function in adults.

### EPHB4 physical interaction with RASA1 is required for the specification and development of LV valves

The N-terminal SH2 domain of RASA1 binds to either of two autophosphorylated tyrosine residues in the EPHB4 juxtamembrane region and thus mediates direct RASA1-EPHB4 physical interaction during the course of EPHB4 signal transduction^11,36^. To assess a requirement of RASA1-EPHB4 physical interaction for vascular development and function, we previously generated an EPHB4 2YP knockin mouse mutant that expresses a form of EPHB4 that is unable to interact with RASA1 physically yet retains protein tyrosine kinase activity^11^. Surprisingly, preliminary genetic analyses indicated no deleterious effects of the mutation in homozygous form, and each of vasculogenesis, developmental blood and lymphatic angiogenesis, neonatal angiogenesis, and pathological angiogenesis in adults were found to proceed normally in *Ephb4^2YP/2YP^* homozygous mice, thus indicating that RASA1-EPHB4 physical interaction is not required for these events ^11^. However, upon analyses of larger numbers of progeny generated from intercrosses of *Ephb4^fl/2YP^* heterozygous parents, we observed a less-than-expected number of *Ephb4^2YP/2YP^* homozygote pups at 2 weeks of age (Figure 3A,B). Significantly, in some instances, we determined that the cause of death prior to weaning was chylothorax, resulting in death by suffocation (Figure 3C). In mice, chylothorax is commonly associated with defective LV valve number and/or function^24,41,42^. Moreover, mice induced to lose expression of EPHB4 in the neonatal period develop chylothorax^9^. Therefore, we examined LV valve numbers in EPHB4 2YP mice. The number of valves in collecting mesenteric LV of P11 *Ephb4^2YP/2YP^*homozygote pups was much reduced compared to littermate *Ephb4^fl/fl^*controls (Figure 3D). Likewise, there were reduced numbers of LV valves in diaphragmatic collecting LV of P11 *Ephb4^2YP/2YP^* homozygotes and in popliteal collecting LV of adults (Figure S2). Analysis of embryonic mesenteric LV valve development showed much-reduced numbers of valves in *Ephb4^2YP/2YP^* homozygotes at E17.5 through E19.5 and impaired LV valve specification at E16.5 (Figure 3E-G). Staining of tissue sections from E19.5 *Ephb4^fl/fl^* and *Ephb4^2YP/2YP^*embryos with an anti-EPHB4 antibody showed that EPHB4 2YP was expressed at similar levels as wild-type EPHB4 in both BEC and LEC (Figure S3). Therefore, impaired LV valve development in *Ephb4^2YP/2YP^*homozygotes cannot be explained by reduced expression of EPHB4 but instead by a required role for RASA1-EPHB4 physical interaction for this event.

**Figure 3.**
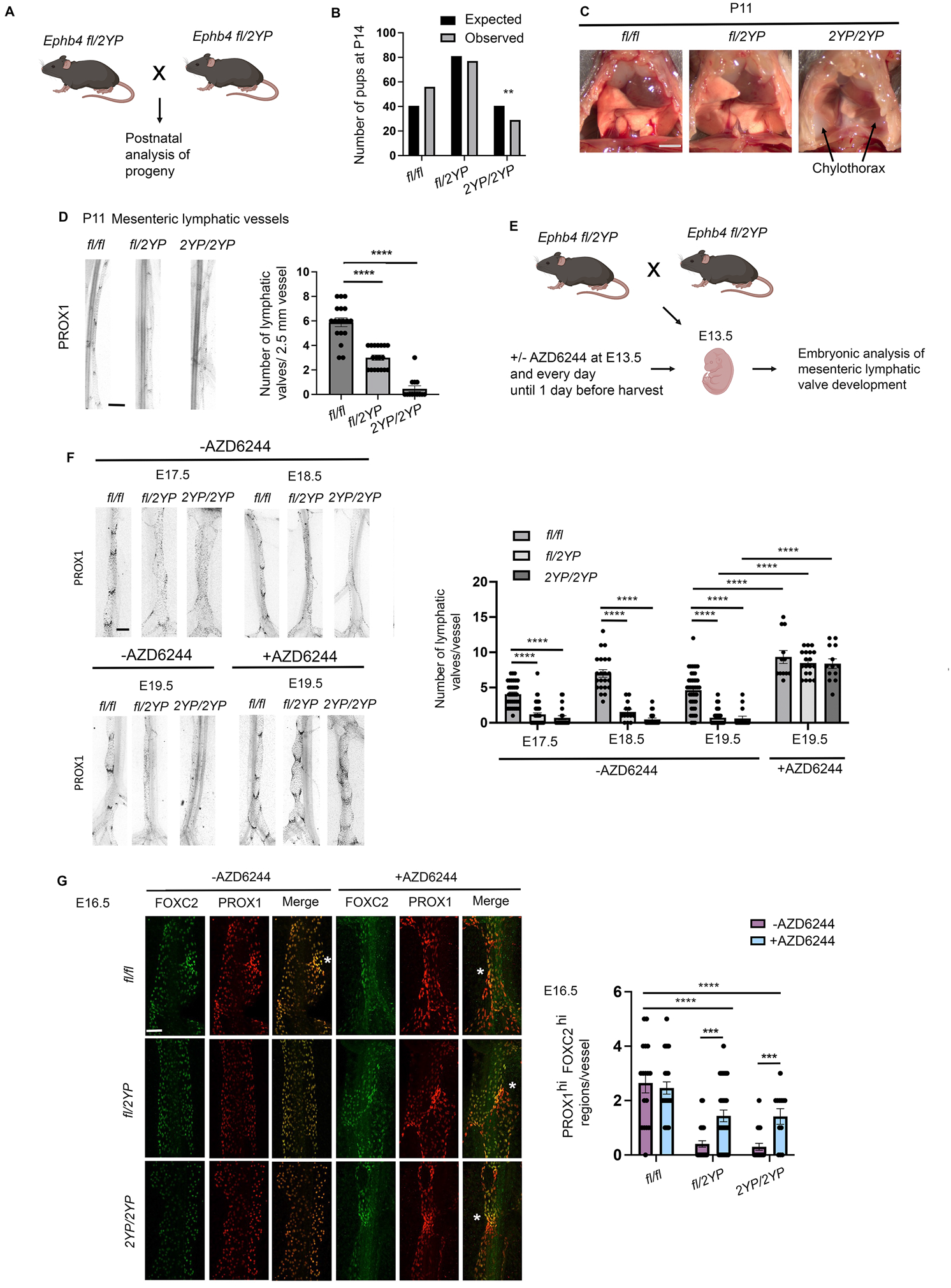
LV valve specification and development in EPHB4 2YP mice. (A) Schematic showing design of experiments in (B-D). (B) Observed and expected numbers of pups of the indicated genotypes at 2 weeks of age. ** *P*<0.01, two-tailed Chi-squared test. (C) Exposed chest cavities of P11 littermate pups showing chylothorax in the *Ephb4 2YP/2YP* homozygote. Scale bar=250 mm. (D) Representative images of mesenteric LV from P11 pups stained with PROX1 are shown at left. Scale bar=200 µm. Graph shows mean +/− 1 SEM of the number of mesenteric LV valves in P11 pups of the indicated genotypes (*Ephb4 fl/fl*, n=18 vessels from 4 mice; *Ephb4 fl/2YP*, n=18 vessels from 5 mice; *Ephb4 2YP/2YP*, n=13 vessels from 3 mice). **** *P*<0.0001, two-tailed Mann-Whitney test. (E) Schematic showing design of experiments in (F and G). (F) Representative images of PROX1-stained mesenteric LV from embryos of the indicated ages and genotypes are shown at left. Scale bar=100 µm. At right is shown the mean +/− 1 SEM of numbers of valves per LV without AZD6244 treatment assessed at E17.5 (*Ephb4 fl/fl*, n=34 vessels from 4 embryos; *Ephb4 fl/2YP*, n=47 vessels from 9 embryos, *Ephb4 2YP/2YP*, n=21 vessels from 4 embryos), E18.5 (*Ephb4 fl/fl*, n=22 vessels from 4 embryos; *Ephb4 fl/2YP*, n=13 vessels from 3 embryos, *Ephb4 2YP/2YP*, n=21vessels from 3 embryos) and E19.5 (*Ephb4 fl/fl*, n=43 vessels from 7 embryos; *Ephb4 fl/2YP*, n=104 vessels from 16 embryos, *Ephb4 2YP/2YP*, n=15 vessels from 3 embryos), and with AZD6244 treatment at E13,5, assessed at E19.5 (*Ephb4 fl/fl*, n=12 vessels from 6 embryos; *Ephb4 fl/2YP*, n=19 vessels from 13 embryos, *Ephb4 2YP/2YP*, n=13 vessels from 4 embryos). **** *P*<0.0001, two-tailed Mann-Whitney test. (G) At left are shown representative images of PROX1 and FOXC2 staining of mesenteric LV. Asterisks in merged images indicate sites of high FOXC2 and PROX1 staining. Scale bar=100 µm. At right is shown the mean +/− 1 SEM PROX1^hi^ FOXC2^hi^ regions per LV without AZD6244 treatment (*Ephb4 fl/fl*, n=17 vessels from 3 embryos; *Ephb4 fl/2YP*, n=32 vessels from 5 embryos, *Ephb4 2YP/2YP*, n=20 vessels from 3 embryos), and with AZD6244 treatment (*Ephb4 fl/fl*, n=28 vessels from 4 embryos; *Ephb4 fl/2YP*, n=32 vessels from 6 embryos, *Ephb4 2YP/2YP*, n=12 vessels from 2 embryos). *** *P*<0.001, **** *P*<0.0001, two-tailed Mann-Whitney test.

Significantly, *Ephb4^fl/2YP^* heterozygotes also showed reduced numbers of valves in P11 mesenteric and diaphragmatic collecting LV, adult popliteal collecting LV, and embryonic mesenteric collecting LV (Figure 3D-F and Figure S2). Furthermore, Ephb*4^fl/2YP^* heterozygotes showed impaired LV valve specification (Figure 3G). In contrast, mice that were heterozygous for an *Ephb4* null allele showed normal LV valve development (Figure S4). These findings indicate that EPHB4 2YP functions in a dominant negative fashion to block LV valve specification and development in mice.

**Figure 4.**
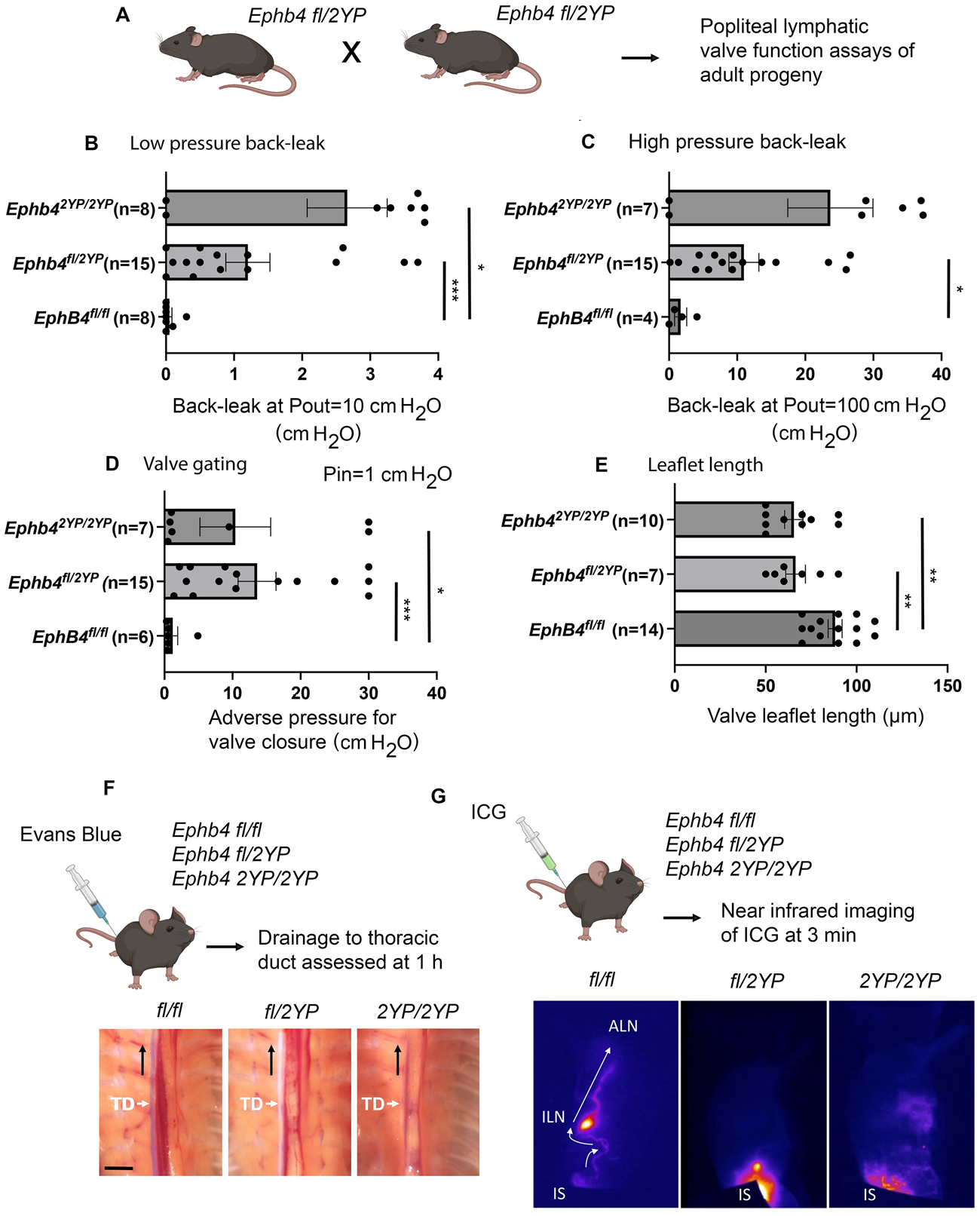
LV valve function in EPHB4 2YP adult mice. (A) Schematic showing design of experiments in (B-E). (B) Mean +/− 1 SEM of the amount of pressure back-leak at a Pout of 10 cm H_2_O. Numbers of tested valves are indicated in parentheses and were obtained from the following numbers of mice: *Ephb4 fl/fl*, n=4; *Ephb4 fl/2YP*, n=5; *Ephb4 2YP/2YP*, n=3. * *P*<0.05, ** *P*<0.01, two-tailed Mann-Whitney test. (C) Mean +/− 1 SEM of the amount of pressure back-leak at a Pout of 100 cm H_2_O. Numbers of tested valves are indicated in parentheses and were obtained from the same number of mice as indicated in (B). * *P*<0.05, two-tailed Mann-Whitney test. (D) Mean +/− 1 SEM of the amount of adverse pressure required to close valves at a Pin of 1 cm H_2_O. Numbers of tested valves are indicated in parentheses and were obtained from the same number of mice as indicated in (B). * *P*<0.05, *** *P*<0.001, two-tailed Mann-Whitney test. (E) Mean +/− 1 SEM of valve leaflet lengths. Numbers of valve leaflets measured are indicated in parentheses and were obtained from the same number of mice as indicated in (B), except for *Ephb4 fl/fl* controls that include *Ephb4 fl/fl* controls treated with TM in Figure 2F (n=9 mice total). ** *P*<0.01, two-tailed Mann-Whitney test. (F) Images show thoracic ducts (TD) in the chest cavity after injection of Evans Blue dye into the base of the tail of P11 mice. Note absence of dye drainage to the TD in *Ephb4 fl/2YP* and *Ephb4 2YP/2YP* mice. Scale bar=2 mm. (G) Near infrared images of indocyanine green (ICG) drainage into the cutaneous/subcutaneous LV network after injection into the base of the tail of adult mice. Note drainage of ICG to the inguinal lymph node (ILN) and toward the axillary lymph node (ALN) in *Ephb4 fl/fl* but not *Ephb4 fl/2YP* and *Ephb4 2YP/2YP* mice. IS, injection site.

To assess if impaired LV valve development in *Ephb4^2YP/2YP^* and *Ephb4^fl/2YP^* mice is consequent to dysregulated Ras-MAPK activation, we administered AZD6244 to embryos at E13.5. AZD6244 rescued LV valve development in both heterozygotes and homozygotes as determined at E19.5 (Figure 3F) and LV valve specification as determined at E16.5 (Figure 3G). Notably, when administered at E13.5, AZD6244 induced larger numbers of valves at E19.5 even in control *Ephb4^fl/fl^* embryos (Figure 3F). This finding indicates that MAPK activation is dispensable for LV valvulogenesis and acts only to inhibit this event.

### Impaired LV valve function in EPHB4 2YP mice

We next analyzed the function of LV valves that developed in EPHB4 2YP mice. The majority of popliteal LV valves from *Ephb4^2YP/2YP^* and *Ephb4^fl/2YP^* mice were dysfunctional in low-pressure back-leak tests and in high-pressure back-leak tests in which Pout is raised to 100 cm H_2_O (Figure 4A-C). In addition, most LV valves from *Ephb4^fl/2YP^* mice were dysfunctional in valve closure tests (Figure 4D). Similar to LV valves from adult induced EPHB4-deficient mice, valve leaflets were on average shorter in *Ephb4^2YP/2YP^* and *Ephb4^fl/2YP^*mice, which accounts for impaired function (Figure 4E). Since *Ephb4^2YP/2YP^*and *Ephb4^fl/2YP^* mice have reduced numbers of LV valves, the majority of which are dysfunctional, we examined if mice showed disordered LV flow. For this purpose, we injected Evans Blue dye subcutaneously into the base of the tail of mice and examined its drainage to the thoracic duct after 1 hour. In contrast to control *Ephb4^fl/fl^* mice, Evans Blue was not detected in the thoracic ducts of *Ephb4^2YP/2YP^*and *Ephb4^fl/2YP^* mice (Figure 4F). In parallel experiments, we also injected indocyanine green intradermally into the base of the tail and monitored its drainage through the cutaneous/subcutaneous LV network by near infrared imaging. In control mice, dye consistently drained to the inguinal lymph node and from that site to the axillary lymph node. In contrast, in *Ephb4^2YP/2YP^* and *Ephb4^fl/2YP^*mice, dye was trapped at the injection site and failed to enter the LV network (Figure 4G). Thus, *Ephb4^2YP/2YP^*and *Ephb4^fl/2YP^* mice show impaired LV flow that is consistent with impaired LV valve development and function and the occurrence of chylothorax.

### Physical interaction between RASA1 and EPHB4 is necessary to limit activation of the Ras-MAPK pathway triggered by oscillatory shear stress

Our findings with EPHB4 2YP mice indicate that physical interaction between RASA1 and EPHB4 is dispensable for developmental BV and LV angiogenesis but necessary for LV valvulogenesis. LV valvulogenesis is thought to be triggered by oscillatory shear stress (OSS) that induces the valvulogenesis differentiation program, including upregulation of expression of FOXC2, PROX1 and the integrin alpha-9 (ITGA9), among other transcriptional changes^39,40,43,44^. Therefore, we investigated if physical interaction between EPHB4 and RASA1 is uniquely required to regulate Ras-MAPK activation triggered in response to OSS. For this purpose, we examined OSS-induced MAPK activation in primary human dermal LEC (HDLEC). To examine if EPHB4 and RASA1 are required to control OSS-induced MAPK activation in HDLEC, cells were treated with siRNA to knockdown the expression of EPHB4 and RASA1 (Figure 5A). In control HDLEC that were treated with a scrambled siRNA, OSS induced a modest transient activation of MAPK that peaked at 10 minutes (Figure 5B). In EPHB4 and RASA1 siRNA-treated HDLEC, increased MAPK activation was observed even under static conditions (Figure 5B). Most likely this reflects roles for EPHB4 and RASA1 as negative regulators of Ras-MAPK activation triggered by LEC-derived growth factors such as VEGF-C (see below). Nonetheless, in both EPHB4- and RASA1-depleted HDLEC, OSS induced robust further activation of MAPK that increased in magnitude through the duration of the experiment up to 60 minutes (Figure 5B). In contrast, a different MAPK activation response was observed when HDLEC were subject to laminar shear stress (LSS) (Figure S5). Thus, although EPHB4 and RASA1 knockdown resulted in increased MAPK activation at 10 minutes, the kinetics of the LSS-induced MAPK activation response was similar in EPHB4 and RASA1 knockdown LEC to that observed in control HDLEC. To determine if physical interaction between RASA1 and EPHB4 is required to limit the OSS-induced MAPK response in HDLEC, we next transduced HDLEC with lentivirus encoding EPHB4 2YP. Since EPHB4 2YP behaves as a dominant negative form of EPHB4 during valvulogenesis in vivo, we reasoned that EPHB4 2YP should behave similarly in HDLEC in vitro and dominate the function of endogenous wild-type EPHB4. Indeed, EPHB4 2YP transduced LEC showed a similar dysregulated OSS-induced MAPK activation response as EPHB4 and RASA1 knockdown HDLEC, with responses persisting through at least 60 minutes (Figure 5B). Thus, physical interaction between EPHB4 and RASA1 is necessary to control the OSS-induced MAPK response in LEC.

**Figure 5.**
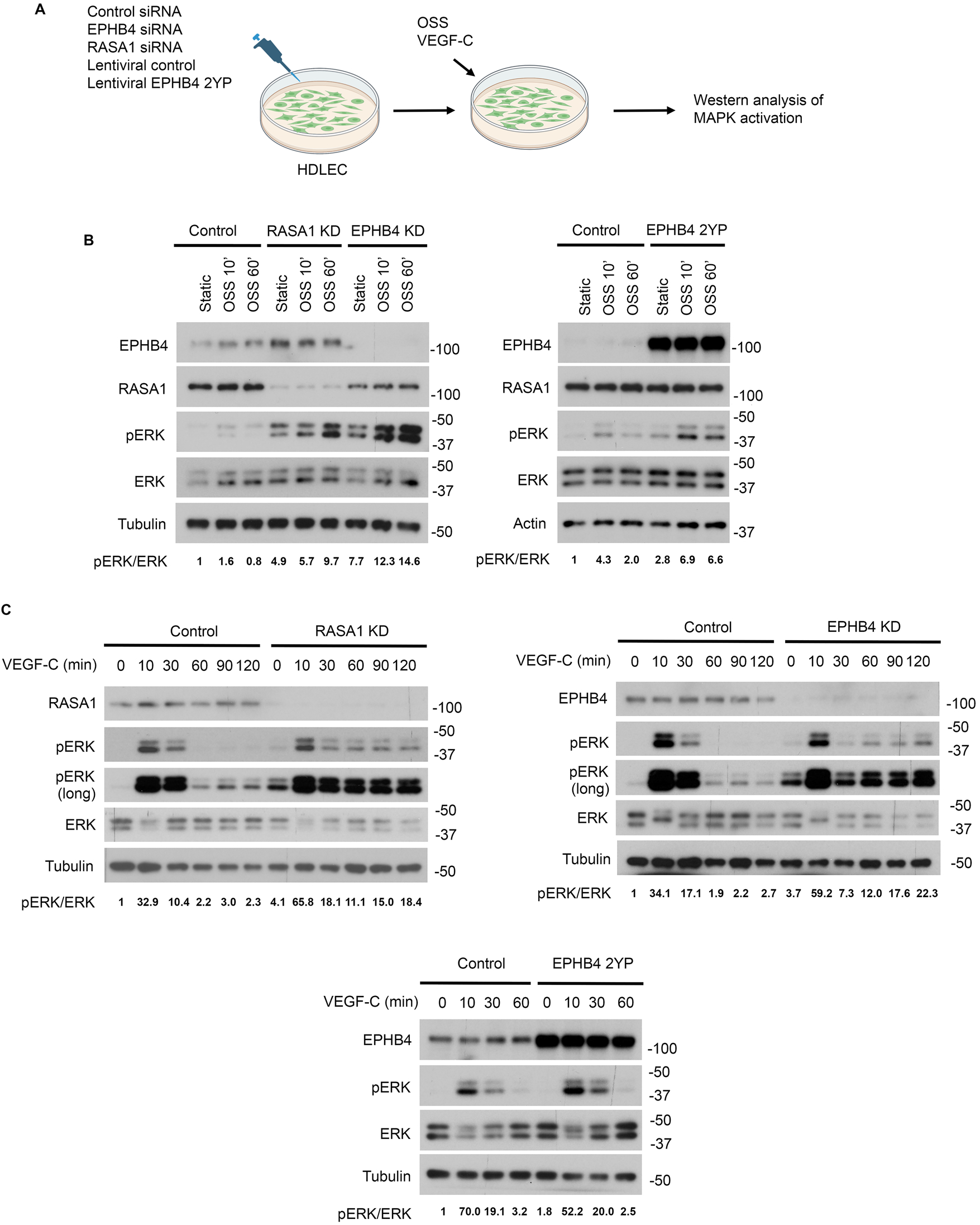
OSS- and VEGF-C-induced activation of MAPK in HDLEC after EPHB4 or RASA1 knockdown or after expression of EPHB4 2YP. (A) Schematic showing design of experiments in (B and C). (B) Left, Western blot analysis of OSS-induced MAPK activation in HDLEC treated with control, RASA1, or EPHB4 siRNA (RASA1 or EPHB4 knockdown, KD). Right, OSS-induced MAPK activation in HDLEC infected with control lentivirus or lentivirus encoding EPHB4 2YP. Blots were probed with anti-tubulin or anti-actin antibodies to demonstrate equivalent protein loading. (C) Western blot analysis of VEGF-C-induced MAPK activation in HDLEC treated with control, RASA1, or EPHB4 siRNA, or control lentivirus or lentivirus encoding EPHB4 2YP. Blots were probed with anti-tubulin antibodies to demonstrate equivalent protein loading.

The finding that developmental lymphangiogenesis proceeds normally in EPHB4 2YP mice predicts that physical interaction between EPHB4 and RASA1 should not be necessary to control growth factor-induced MAPK activation in LEC. To address this question, we examined MAPK activation in HDLEC triggered by VEGF-C, which is the principal growth factor that mediates lymphangiogenesis^45^. We demonstrated previously that lung LEC derived from induced adult RASA1-deficient mice show persistent MAPK activation after stimulation with VEGF-C^27^. In agreement with this, HDLEC treated with RASA1 siRNA showed persistent activation of MAPK in response to VEGF-C (Figure 5C). Likewise, HDLEC treated with EPHB4 siRNA showed persistent activation of MAPK in response to VEGF-C (Figure 5C). These findings are consistent with our previous discoveries of disrupted developmental lymphangiogenesis in induced RASA1- and EPHB4-deficient embryos that can be rescued by blockade of MAPK signaling^11,26^. However, in stark contrast to OSS-induced MAPK activation, VEGF-C-induced MAPK activation did not persist in HDLEC transduced with EPHB4 2YP (Figure 5C). Thus, physical interaction between EPHB4 and RASA1 is necessary to control OSS-induced MAPK activation but not VEGF-C-induced MAPK activation, again consistent with in vivo findings.

### An EPHB4-RASA1 signaling complex is necessary for OSS-induced induction of LV valve leaflet markers

In agreement with the concept that OSS is the trigger for LV valvulogenesis in vivo, LEC subjected to OSS in vitro upregulate LV valve leaflet markers^43,44^. Since physical interaction between RASA1 and EPHB4 is necessary for LV valvulogenesis in vivo, we reasoned that physical interaction should also be necessary for OSS-induced upregulation of LV valve markers in HDLEC in vitro. Therefore, we examined if knockdown of EPHB4 or RASA1 or expression of EPHB 2YP influenced OSS-induced expression of ITGA9 and FOXC2 in HDLEC (Figure 6A). In control HDLEC, OSS consistently increased the expression of both valve markers. In contrast, each of EPHB4 knockdown, RASA1 knockdown, and expression of EPHB4 2YP blocked OSS-induced induction of ITGA9 and FOXC2 (Figure 6B). Thus, RASA1-EPHB4 physical interaction is necessary for OSS-induced expression of valve markers in HDLEC in vitro, consistent with findings in vivo.

**Figure 6.**
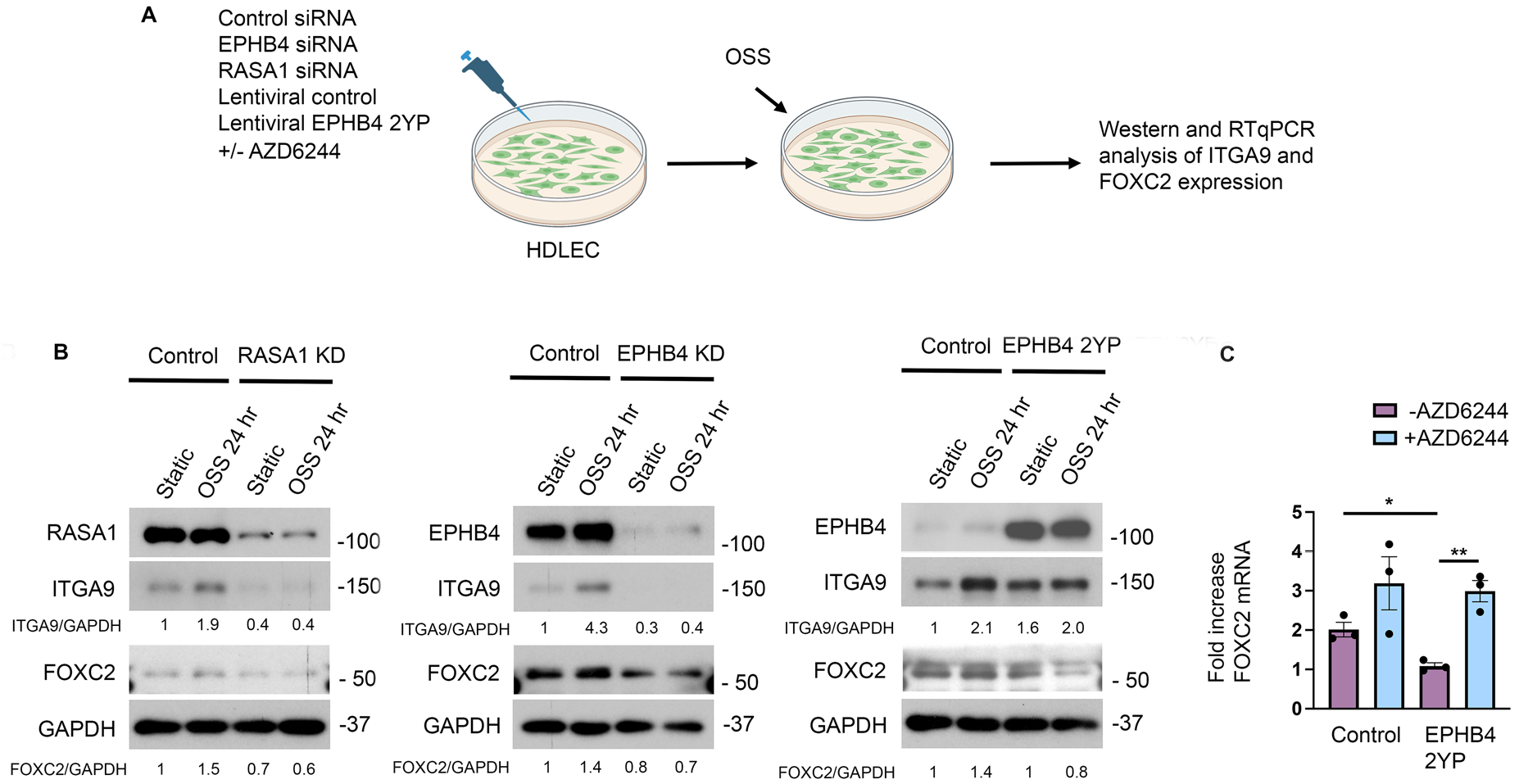
OSS-induced expression of ITGA9 and FOXC2 in HDLEC after EPHB4 or RASA1 knockdown or after expression of EPHB4 2YP. (A) Schematic showing design of experiments in (B and C). (B) Western blot analysis of OSS-induced expression of ITGA9 and FOXC2 in HDLEC treated with control, RASA1, or EPHB4 siRNA, or control lentivirus or lentivirus encoding EPHB4 2YP. Blots were probed with anti-GAPDH antibodies to demonstrate equivalent protein loading. (C) Quantitative RT-PCR analysis of *FOXC2* mRNA expression in HDLEC transduced with control lentivirus or lentivirus encoding EPHB4 2YP subject to OSS for 48 hours in the presence or absence of AZD6244 (n=3 each condition). Fold increases are relative to HDLEC under static conditions without AZD6244 for each HDLEC type. * *P*<0.05, ** *P*<0.01, two-tailed Student’s t-test.

Inhibition of MAPK signaling rescues impaired LV valve specification and development in EPHB4 2YP embryos (Figure 3). Based on these findings, we predicted that inhibition of the MAPK pathway should also rescue OSS-induced upregulation of LV valve markers in EPHB4 2YP-expressing HDLEC. For FOXC2, we examined this at the transcriptional level by quantitative RT-PCR to detect *FOXC2* mRNA transcripts (Figure 6A). OSS consistently upregulated *FOXC2* mRNA levels in control HDLEC. In contrast, OSS did not upregulate

*FOXC2* mRNA in HDLEC transduced with EPHB4 2YP (Figure 6C). However, the inclusion of AZD6244 into the medium for the duration of exposure to OSS rescued the induction of *FOXC2* mRNA expression, consistent with in vivo findings and further supporting the view that in the absence of EPHB4-RASA1 physical interaction, dysregulated Ras-MAPK signaling impedes LV valve specification and development (Figure 6C).

### Physical interaction between EPHB4 and RASA1 is necessary to control MAPK activation induced by the Piezo1 OSS sensor and for Piezo-1-induced LV valvulogenesis in vivo

Piezo1 is a mechanosensor that is necessary for LV valvulogenesis in vivo and for OSS-induced increased valve marker expression in HDLEC in vitro^46,47^. In addition, Piezo1 is known to trigger activation of the Ras-MAPK pathway in HDLEC and other cell types^46,48^. Therefore, we considered that a requirement for physical interaction between RASA1 and EPHB4 in the regulation of OSS-induced MAPK activation in HDLEC might be explained by a required role for physical interaction in the regulation of Piezo1-induced MAPK activation. To address this, we examined MAPK activation in HDLEC induced by the Piezo1 agonist, Yoda-1^49^ (Figure 7A). Consistent with previous reports, Yoda-1 induced a transient activation of MAPK in HDLEC that peaked around 10 min^46,48^. However, after knockdown of EPHB4 or RASA1 or after expression of EPHB4 2YP in HDLEC, Yoda1-induced MAPK activation persisted through at least 90 minutes (Figure 7B). Therefore, physical interaction between RASA1 and EPHB4 is necessary to limit Piezo1-induced MAPK activation in LEC.

**Figure 7.**
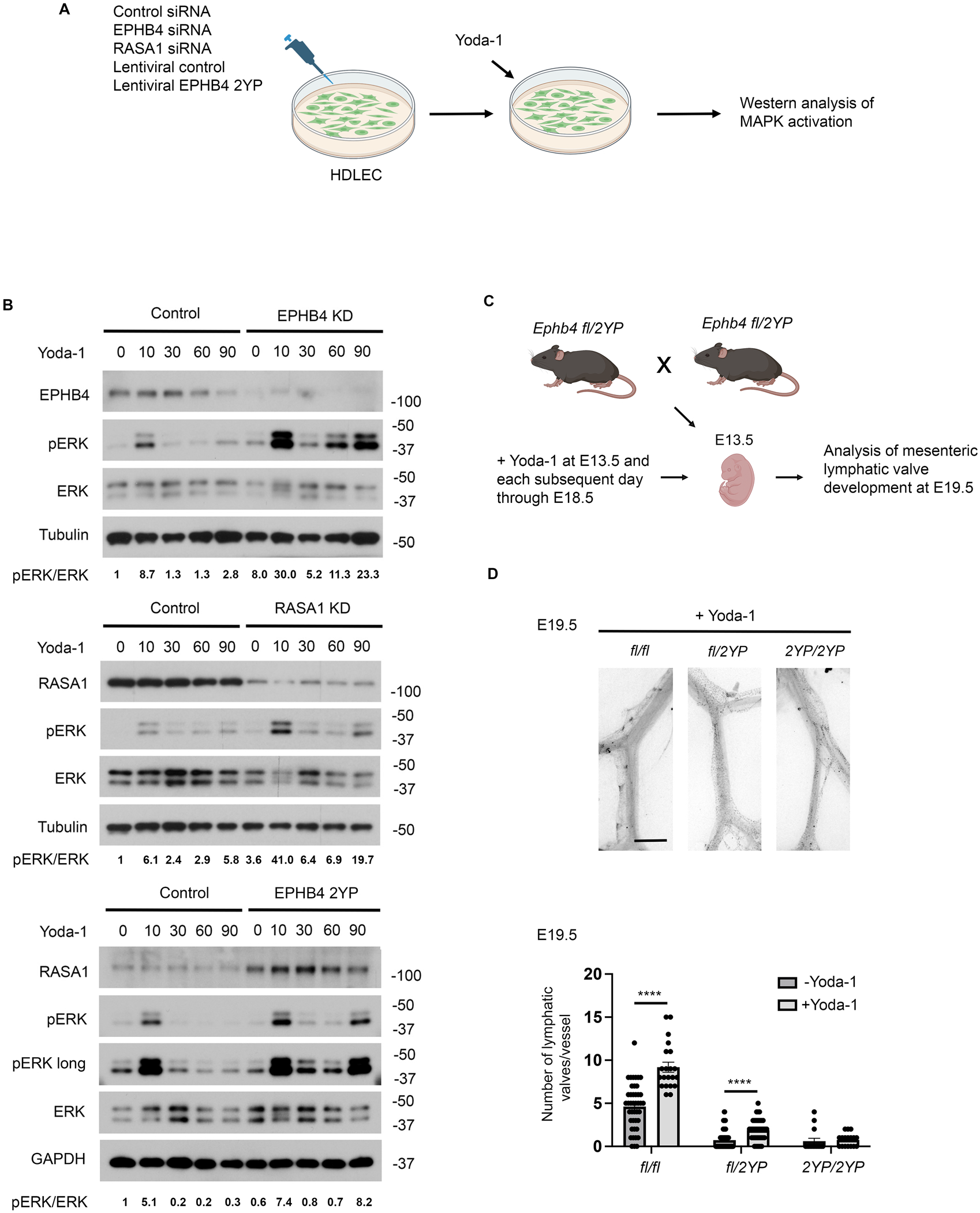
EPHB4 and RASA1 regulation of Piezo1-induced MAPK activation in HDLEC and Piezo-1-induced LV valvulogenesis in vivo. (A) Schematic showing design of experiments in (B and C). (B) Yoda-1-induced MAPK activation in HDLEC treated with control, RASA1, or EPHB4 siRNA, or control lentivirus or lentivirus encoding EPHB4 2YP. Blots were probed with anti-tubulin or anti-GAPDH antibodies to demonstrate equivalent protein loading. (C) Schematic showing design of experiments in (D). (D) Images of mesenteric LV from E19.5 embryos stained with PROX1 are shown at top. Scale bar=200 µm. Graph below shows mean +/− 1 SEM of the number of mesenteric LV valves in E19.5 embryos of the indicated genotypes treated with Yoda-1 (*Ephb4 fl/fl*, n=21 vessels from 4 embryos; *Ephb4 fl/2YP*, n=41 vessels from 8 embryos; *Ephb4 2YP/2YP*, n=17 vessels from 3 embryos). Comparisons are made with E19.5 embryos not treated with AZD6244 from Figure 3F (*Ephb4 fl/fl*, n=43 vessels from 7 embryos; *Ephb4 fl/2YP*, n=104 vessels from 16 embryos, *Ephb4 2YP/2YP*, n=15 vessels from 3 embryos). **** *P*<0.0001, two-tailed Mann-Whitney test.

To confirm that physical interaction between EPHB4 and RASA1 is necessary to control Piezo-1-induced LV valvulogenesis in vivo, we examined the effect of Yoda-1 upon valvulogenesis in EPHB4 2YP mice. For this purpose, we administered Yoda-1 to E13.5 embryos from crosses of *Ephb4^fl/2YP^* heterozygotes and assessed mesenteric LV valve number at E19.5 (Figure 7C). It has been shown previously that Yoda-1 increases mesenteric LV valve number when administered to wild-type embryos in mid to late gestation^46^. In agreement with these findings, Yoda-1 increased mesenteric LV valve number in control *Ephb4^fl/fl^* embryos. However, Yoda-1 was much less effective at inducing increased numbers of LV valves in *Ephb4^fl/2YP^* embryos and was completely unable to increase LV valve number in *Ephb4^2YP/2YP^* homozygote embryos (Figure 7D). These findings support the contention that physical interaction between EPHB4 and RASA1 is necessary to regulate Piezo-1-induced LV valvulogenesis.

To determine if physical interaction between EPHB4 and RASA1 is also required to regulate MAPK activation induced by additional OSS sensors in HDLEC, we examined responses in the presence of the Piezo1 inhibitor, GsTMx4^50^ (Figure S6A). Consistent with the ability of Yoda1 to activate MAPK in HDLEC at 10 min, GsMT4 blocked OSS-induced MAPK activation at this time point. Instead, a modest later activation of MAPK was observed at 60 minutes in control cells, indicating the participation of additional OSS sensors in the activation of MAPK under these conditions (Figure 6B). Notably, this later activation of MAPK in the presence of GsMTx4 was substantially increased in HDLEC transduced with EPHB4 2YP (Figure S6B). Therefore, EPHB4-RASA1 physical interaction may also be required to limit MAPK activation triggered through other undefined OSS sensors as well as Piezo1.

## DISCUSSION

Evidence is emerging that a major function of EPHB4 in EC in vivo is to limit the activation of the Ras-MAPK pathway. This was initially demonstrated for developmental angiogenesis, during which disruption of EPHB4 in EC led to dysregulated activation of the Ras-MAPK pathway followed by EC apoptotic death that could be rescued by drugs that inhibit this pathway^6,26^. In this study, we now demonstrate that EPHB4-mediated inhibition of Ras-MAPK signaling is essential for the specification of lymphatic vessel valves and for their continued development after specification. The finding that dysregulated Ras-MAPK activation in the absence of EPHB4 inhibits LV valvulogenesis contrasts with the findings of an earlier study in which it was shown that dysregulated Ras-MAPK activation promoted the differentiation of LEC from cardinal vein EC during the process of lymphatic vessel specification through upregulation of SOX18 and PROX1 transcription factor expression^51^. Ras-MAPK-mediated inhibition of LV specification involves suppression of the expression of the FOXC2 transcription factor that acts upstream of PROX1 in valvulogenesis^44,52^. However, whether MAPK inhibits FOXC2 expression directly or indirectly through inhibiting transcription factors upstream of FOXC2, such as GATA2^53^, remains to be determined.

Concerning the mechanism by which dysregulated Ras-MAPK activation in the absence of EPHB4 impairs continued lymphatic valve development after specification, this is most likely explained by the retention of collagen IV in LV valve leaflet EC during valve elongation. LV leaflet elongation requires LEC synthesis of extracellular matrix proteins, including collagen IV, that are normally deposited in the extracellular matrix core of the valve leaflet^39,40^. Consequently, EPHB4-deficient LEC would be expected to undergo apoptosis during valve elongation due to an unresolved UPR or alternatively would detach from leaflets and meet their ultimate demise in the vessel lumen as a result of anoikis. Dysregulated Ras-MAPK activation in EC in the absence of RASA1 blocks developmental angiogenesis through this mechanism that is also responsible for the impaired development of lymphovenous valves and venous valves^25,26^. In addition, a similar mechanism may account for the role of EPHB4 in lymphatic valve maintenance in adults. Subject to high shear stress forces, LEC in valve leaflets may need to continue to engage in high-rate collagen IV synthesis to remain attached to the leaflet core. Indeed, shear stress is known to upregulate EC synthesis of collagen IV in vitro^54^. Thus, in the absence of EPHB4, collagen IV would become trapped within the endoplasmic reticulum of established valve leaflet LEC leading to their apoptosis and detachment from leaflets. This would be consistent with the loss of LEC in valve leaflets in adult induced EPHB4-deficient mice. However, whether the role of EPHB4 in LV valve maintenance relates to its function as an inhibitor of Ras-MAPK signaling is not yet certain, given that AZD6244 caused LV valve dysfunction in adult wild-type mice.

The JM region of EPHB2, which is highly homologous to EPHB4 in its intracellular region, folds as an alpha-helix that interacts with the N and C lobes of the EPHB4 kinase domain to restrain EPHB2 kinase activity^55,56^. During the course of EPHB2 signal transduction, autophosphorylation of two tyrosine residues in the JM region disrupts this helical structure and switches the kinase into an open active conformation^55,56^. The RASA1 N-terminal SH2 domain interacts physically with these same two phosphorylated tyrosine residues and thus mediates EPHB4-RASA1 association^36^. With these considerations, simple mutation of these two tyrosine residues to phenylalanine within EPHB4 would not allow conclusions to be readily drawn regarding a requirement of EPHB4-RASA1 physical association for biological effects since mutation of these residues alone would prevent EPHB4 from switching to its fully active state. Therefore, to address this, we previously generated an EPHB4 2YP knockin mouse strain that expresses a form of EPHB4 that contains additional mutations of proline residues in the +3 position of each tyrosine, to glycine^11^. These proline to glycine mutations are predicted to disrupt the alpha-helical conformation of the JM region independently of JM tyrosine phosphorylation, thus obviating a requirement of JM tyrosine phosphorylation for EPHB4 to switch to the active state^55,56^. As expected, EPHB4 2YP cannot interact physically with RASA1 yet retains PTK activity^11^. Surprisingly, EPHB4 2YP mutant mice showed normal blood and lymphatic developmental angiogenesis, indicating that physical interaction between EPHB4 and RASA1 is not required for these events. In contrast, in the current study, we show that LV valvulogenesis depends upon EPHB4-RASA1 physical association. Since physical association between EPHB4 and RASA1 does not result in the modulation of enzymatic activity of either protein^36^, it is likely that the purpose of physical interaction is to target RASA1 to the plasma membrane allowing its juxtaposition to Ras-GTP.

It is intriguing that there is a specific requirement for EPHB4-RASA1 physical interaction for LV valvulogenesis but not lymphatic developmental angiogenesis. As shown in this study, this requirement reflects a need for EPHB4-RASA1 physical interaction to control MAPK activation in LEC induced by OSS but not by growth factors such as VEGF-C. We further show that EPHB4-RASA1 physical association is required to limit MAPK activation in LEC triggered through the Piezo1 OSS sensor, which is essential for LV valvulogenesis^46,47^. This finding could potentially explain a requirement for EPHB4-RASA1 physical interaction for valvulogenesis. In this regard, in a recent study, it was reported that Piezo1 is primarily located in cholesterol-rich caveolae microdomains in the plasma membrane of EC^57^. Therefore, Ras-GTP generated as a result of Piezo1 sensing of OSS is likely to be concentrated in caveolae, at least initially, necessitating that RASA1 be targeted to caveolae to control Piezo1-induced Ras activation. EphB family members, including EPHB4, are also known to localize to caveolae^12^. Thus, one way in which RASA1 could be targeted to caveolae is through physical interaction with EPHB4. However, Piezo1 may not be the only OSS sensor in LEC that is regulated by EPHB4-RASA1 physical interaction, since dysregulated Ras-MAPK activation in response to OSS was also observed in EPHB4 2YP LEC in the presence of a Piezo1 antagonist (Figure S6).

Piezo1 is recognized to function as a mechanosensitive cation channel in multiple cell types^58^. Therefore, how Piezo1 activates the Ras-MAPK pathway in LEC is not immediately obvious. One possibility is that Piezo1-induced intracellular calcium ion influx activates Ras guanine nucleotide exchange factors in LEC that convert Ras from its inactive GDP-bound to its active GTP-bound state^59^. Candidates include Ras guanine nucleotide-releasing proteins (Rasgrps) that contain calcium-responsive EF-hand loops^60–62^. Alternatively, it is possible that intracellular calcium inhibits the GAP activity of one or more RasGAPs, including RASA1 itself, through interaction with RasGAP calcium-binding C2 domains^21^. Regardless of the mechanism, however, any amount of Ras-MAPK activation in LEC inhibits LV development. This can be interpreted from the finding that the administration of AZD6244 to E13.5 embryos substantially increased LV valve number in wild-type embryos as assessed at E19.5 (Figure 3F). Why Piezo1 should, on the one hand, activate signaling pathways that promote valvulogenesis while, on the other hand, activate the Ras-MAPK pathway that opposes this program is uncertain. However, as noted above, Ra-MAPK activation is necessary for LV valve maintenance. Therefore, the dampening of Ras-MAPK signaling through an EPHB4-RASA1 physical interaction mechanism during valvulogenesis may represent a compromise that permits valvulogenesis in the embryo yet allows valve maintenance in adults.

Our results do not formally exclude the possibility that the loss of binding of an SH2 domain-containing protein other than RASA1 to EPHB4 is responsible for the LV valve phenotypes in EPHB4 2YP mice. The Nck adapter protein, for example, has been shown to interact with tyrosine phosphorylated EPHB2, albeit that direct interaction of Nck with EPHB2 has not been demonstrated^63^. However, that the loss of binding of RASA1 in the EPHB4 2YP mutant is the critical factor is supported by the finding that disruption of *Ephb4* and *Rasa1* genes in mice results in similar LV valve phenotypes associated with dysregulated Ras-MAPK signaling in LEC in vivo^24,25^ (and this study). Furthermore, disruption of *EPHB4* and *RASA1* gene expression in human LEC in vitro results in dysregulated Ras-MAPK signaling in response to OSS and Piezo1 stimulation (this study). One other caveat of our study is that although EPHB4-RASA1 physical interaction is shown to be necessary for LV valve specification, the severe impact of the EPHB4 2YP mutation upon valve specification, does not permit conclusions to be drawn regarding a requirement of physical interaction for continued LV valve development or maintenance.

## METHODS

### KEY RESOURCES TABLE

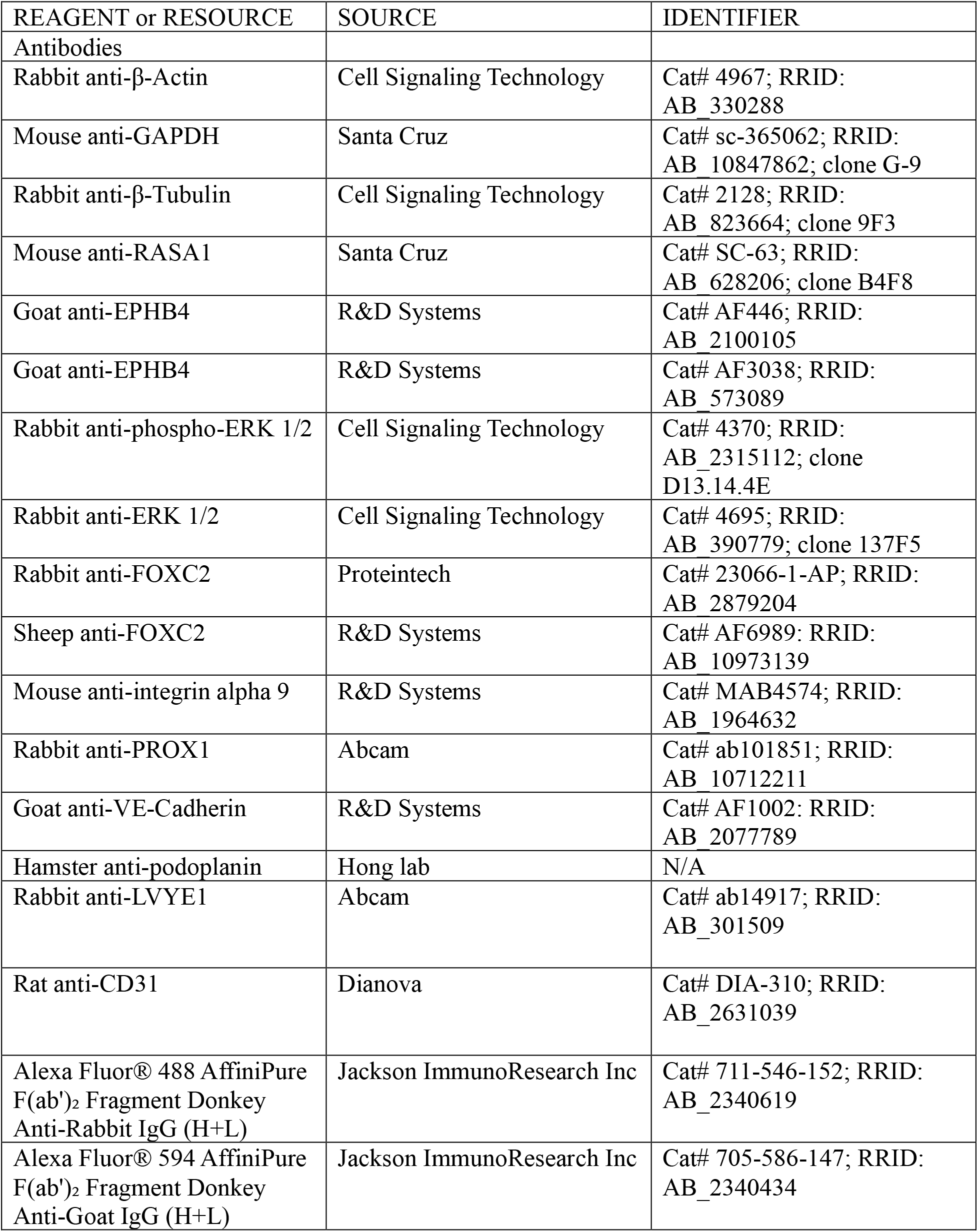
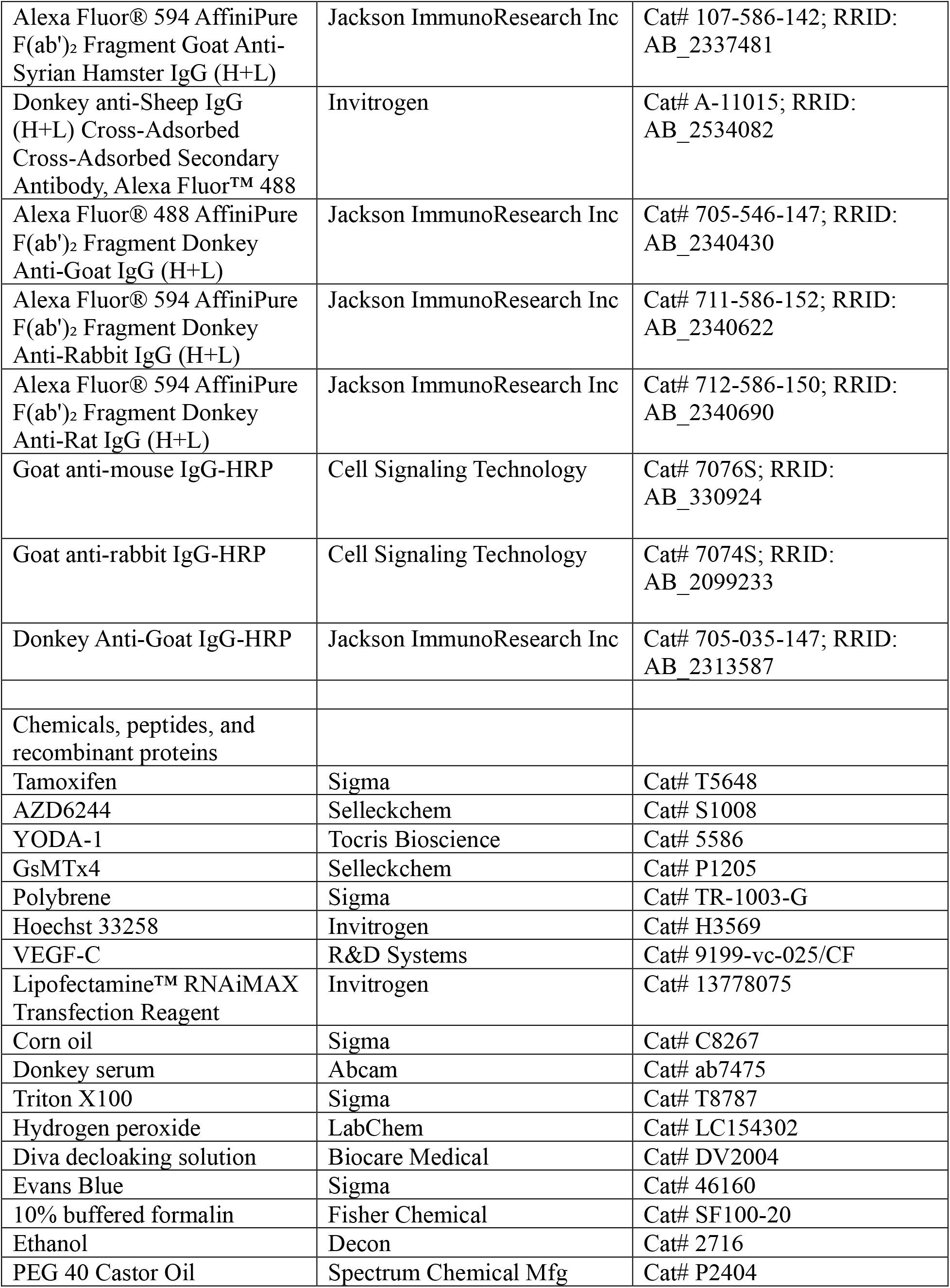
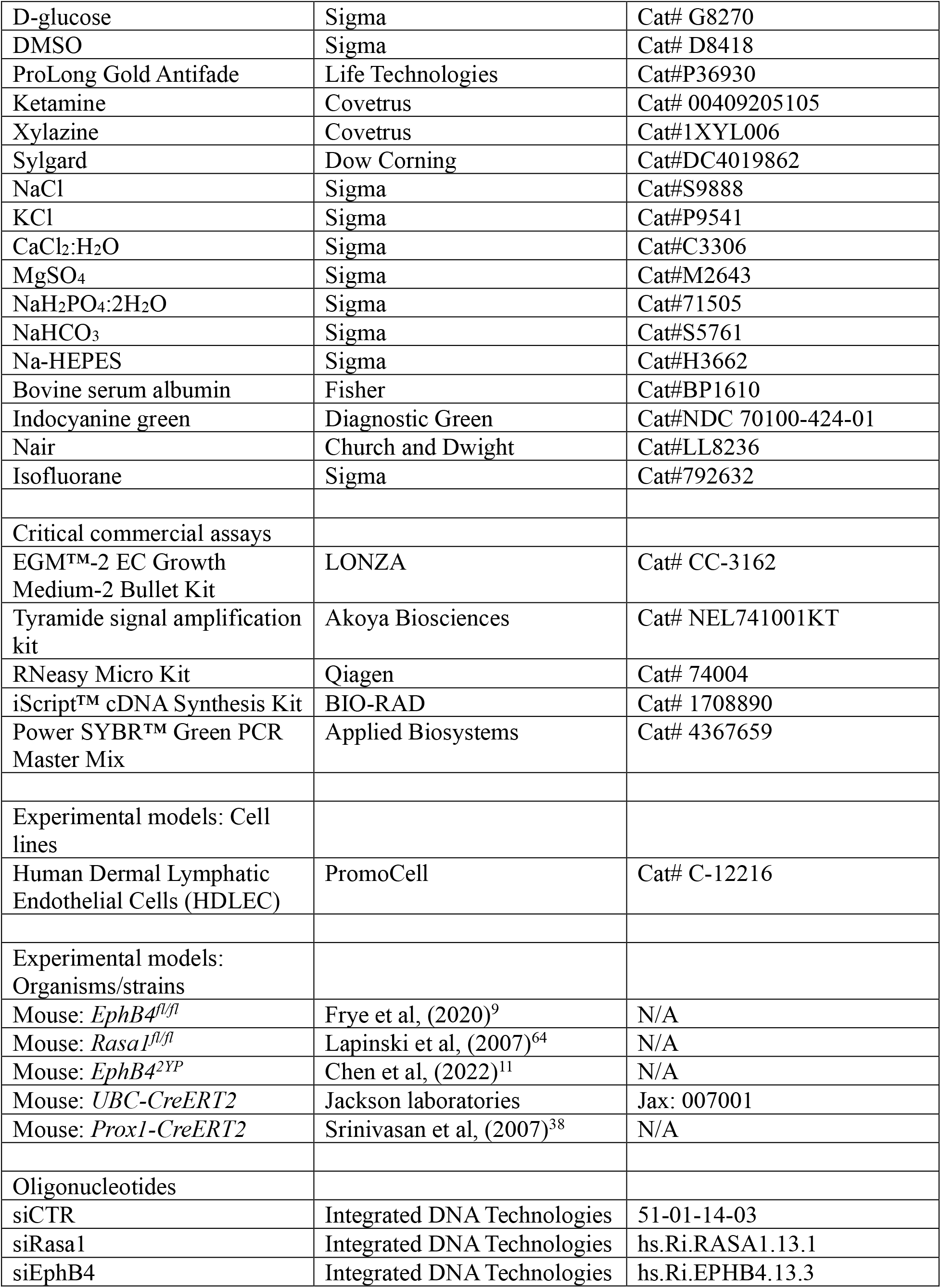
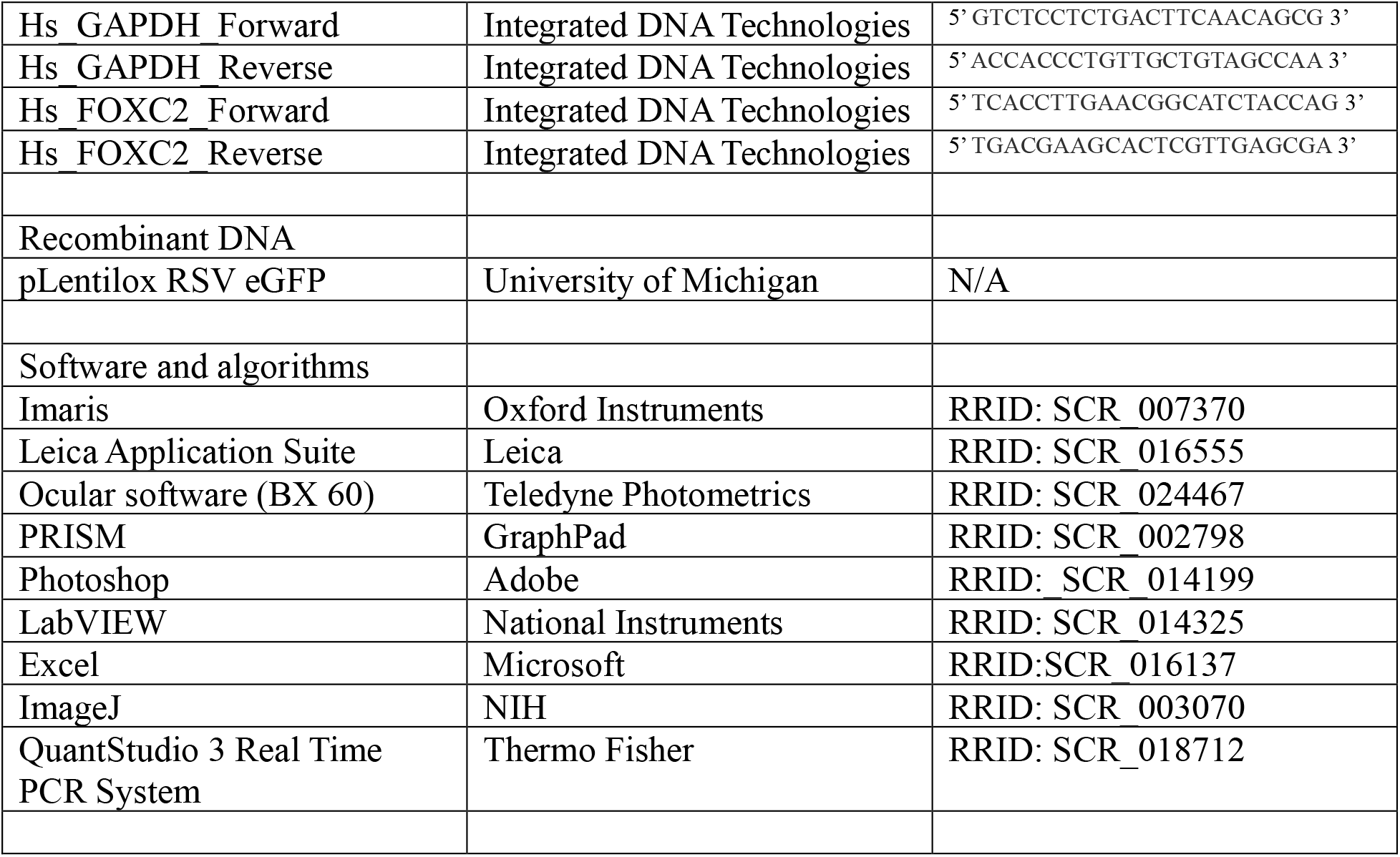

### Animals

All experiments performed with mice complied with University of Michigan, University of Missouri, and University of Texas guidelines and were approved by the respective university committees on the use and care of animals. Mice were on mixed C57BL/6J, 129S6/SvEv and SJL backgrounds. Mice of both sexes were used in studies. No influence of gender upon experimental phenotypes was noted. Where possible, littermate mice and embryos were used as controls.

### Cell line

HDLEC were grown on culture dishes coated with 0.1% gelatin and were maintained in EGM-2 EC Growth Medium-2. Experiments were conducted using cells that were passaged fewer than 8 times.

### Drug administration to mice

Tamoxifen (dissolved in corn oil), AZD6244 (dissolved in 10% Ethanol, 10% PEG40 CASTOR OIL and 80% 5% D-glucose in PBS, and injected 1 mg in each day), and Yoda-1 (50 μg/day in DMSO/ml) were administered to pregnant dams, postnatal mice, and non-pregnant adult mice by i.p. injection at the times and frequencies indicated in the figures. The tamoxifen dose was 1mg/day for pregnant dams and non-pregnant adults and 0.2 mg/day for postnatal mice.

### LV valve enumeration and structural analysis

Mesenteries and diaphragms were harvested and fixed in 1% PFA in PBS overnight. Popliteal collecting LV with associated connective tissue were harvested and fixed in 4% PFA in PBS overnight. Tissues were washed in PBS and blocked in PBS/0.3% Triton-X100/5% donkey serum overnight, and then stained with rabbit anti-PROX1, sheep anti-FOXC2, and goat anti-VE-Cadherin antibodies (1:100 dilution) overnight. After washing in PBS, tissues were incubated with species-specific anti-immunoglobulin donkey F(ab)_2_ fragments coupled to Alexa Flour 488 or 594 overnight (1:500 dilution) followed by further washing in PBS, mounting in ProLong Gold Antifade and viewing on BX60 fluorescence and Nikon A1R confocal microscopes (Nikon).

### Phospho-MAPK immunostaining staining of popliteal LV sections

Popliteal collecting LV fixed as above, were embedded in paraffin. Five-micrometer tissue sections were deparaffinized in xylene, rehydrated in ethanol, incubated for 30 minutes in 3% H_2_O_2_, and boiled in Diva decloaking solution for 30 minutes for antigen retrieval. After blocking in PBS containing 0.1% Triton X-100 and 10% donkey serum for 1 hour, sections were stained with rabbit anti-phospho-ERK (1:100 dilution), and signals were detected with the use of a tyramide signal amplification kit according to the manufacturer’s instructions. Subsequently, sections were stained with a hamster anti-podoplanin antibody (8.1.1, provided by Y. Hong, University of Southern California, Los Angeles, California, USA) overnight, washed and stained with anti-hamster immunoglobulin donkey F(ab)_2_ fragments coupled to Alexa Flour 594 (1:500 dilution) for 1 hour. Slides were washed, mounted, and viewed as above.

### Embryo section immunostaining

Embryos were fixed in 10% buffered formalin overnight, and 5 µm tissue paraffin-embedded sections were treated with Diva decloaker and blocked as above. Sections were stained with goat anti-EPHB4 (AF466), rat anti-CD31, and rabbit anti-LYVE-1 antibodies (1:100 dilution) overnight, washed, and stained with species-specific anti-immunoglobulin donkey F(ab)2 fragments coupled to Alexa Flour 488 or 594 for 1 hour. Sections were washed, mounted, and viewed as above.

#### Popliteal LV isolation, cannulation and pressure control

Mice were anesthetized with ketamine/xylazine (100/10 mg/kg, i.p.) and placed face up on a heated tissue dissection pad. Popliteal lymphatic vessels were exposed through a superficial incision in the leg, removed and transferred to a Sylgard-coated dissection chamber filled with Krebs-albumin solution. A vessel was cleaned of fat and connective tissue, then transferred to a pressure myography chamber where it was cannulated at each end with a glass micropipette (40-50 mm outer diameter tip). After shortening the segment to a single valve, the chamber was moved to the stage of an inverted microscope. The back of each micropipette was connected to a low-pressure transducer and a computerized pressure controller, allowing independent control of inflow (Pin) and outflow (Pout) pressures, as described previously^65^. Custom LabVIEW programs acquired analog data from the pressure transducers simultaneously with vessel inner diameter^66^. Valve test protocols were performed in Ca^2+^-free Krebs buffer at 37°C to prevent pressure fluctuations that otherwise could have interfered with valve function tests.

#### LV valve function tests

A servo-null micropipette (tip diameter = 3-5 mm) was inserted through the wall of the vessel on the inflow side of the valve to measure luminal pressure (Psn). The Psn signal was then calibrated by adjusting the gain and offset of the amplifier/computer while raising Pin and Pout simultaneously between 0.5 and 10 cmH_2_O. To measure back leak through a closed valve, Pin and Pout were set to a relatively low pressure (0.5 cmH_2_O) which set the vessel diameter at ~40% of maximal diameter to facilitate valve closure^65^. Pout was raised to 10 cm H_2_O, ramp-wise, over a 35-second period while Pin was held at 0.5 cmH_2_O. Normal valves closed as Pout exceeded ~1 cmH_2_O and remained closed for the duration of the Pout ramp. Pressure back leak through the closed valve was measured with the Psn micropipette, which could resolve changes as small as ~0.05 cmH_2_O on the inflow side of the valve. This test was repeated 3 times and the values of back leak (Psn – Pin) were averaged to obtain a single representative point for each valve.

In some of our previous studies^67^, we found that only subtle changes in valve function are detectable when the outflow pressure is raised to 10 cmH_2_O. Since normal lymphatic valves can resist much higher pressures (up to 100 cmH_2_O), we also implemented a high-pressure back leak test, in which differences in back leak between normal and dysfunctional valves are often amplified. In this test, the Pin is held at 0.5 cmH_2_O and outflow pressure is raised from 0.5 to 10 cmH_2_O and then to 100 cm H_2_O in steps of 10 cmH_2_O. Psn is measured on the inflow side of the valve and a moveable reservoir is used to raise outflow pressure, otherwise the safe limit of the pressure sensors would be exceeded. Back leak (Psn – Pin) is then plotted as a function of Pout from 0.5 to 100 cmH_2_O.

The third test determined the adverse pressure gradient (ΔP, Pout - Pin) required to close an initially open valve. As demonstrated previously^24,68^, this value increased with increasing vessel diameter. The measurements, therefore, were made at two different baseline pressures (1 or 10 cmH_2_O), which set baseline diameter at ~50% or 100% of maximum diameter. Starting with the valve open, output pressure was raised ramp-wise and the ΔP was determined at the instant of valve closure. The highest ΔP that could be tested was 30 cmH_2_O (equating to a maximum Pout of 40 cm H_2_O when Pin was 10 cm H_2_O) without exceeding the specified safe range of the pressure sensor elements. The data from both valve tests were compiled in Excel and then plotted and analyzed in Prism.

#### Solutions and Chemicals used in LV function tests

Krebs buffer contained: 146.9 mM NaCl, 4.7 mM KCl, 2 mM CaCl_2_·2H_2_O, 1.2 mM MgSO_4_, 1.2 mM NaH_2_PO_4_·H_2_O, 3 mM NaHCO_3_, 1.5 mM Na-HEPES, and 5 mM D-glucose (pH = 7.4). A buffer of the same composition (“Krebs-albumin”) also contained 0.5% bovine serum albumin. Krebs-albumin solution was present both luminally and abluminally during vessel cannulation, and the abluminal solution was constantly exchanged with Ca^2+^-free Krebs solution during the valve function protocols. For Ca^2+^-free Krebs, 3 mM EGTA replaced CaCl_2_·2H_2_O. Purified BSA was obtained from (US Biochemicals; Cleveland, OH), all other chemicals were obtained from Sigma.

### Evans Blue injection

Evans blue (5 mg/ml in PBS) was injected s.c. into hind foot pads and the base of the tail (50 μl at each injection site). After 1 hour, mice were euthanized, and drainage of dye to the thoracic duct was examined.

### Near-infrared imaging of lymphatic flow

A 165.5 µM solution of Indocyanine green (ICG) was prepared in 0.9% NaCl. Mice were shaved from the base of the tail to just under the clavicle, and the remaining hair was removed with Nair™. Mice were anesthetized using isoflurane and intradermally injected with ICG dosing solution in the skin adjacent to the base of the tail. The animals were then illuminated with a 785 nm NIR laser. After masking the site of injection to prevent oversaturation of the image, fluorescent light emission signals were collected by an intensified, cooled 16 bit charged-coupled device with an image integration time of 200 ms as previously described^27^.

### siRNA transfection and viral transduction of HDLEC

HDLEC were seeded into wells of 6 well plates and transfected with (50nM) control, RASA1, or EPHB4 siRNAs using Lipofectamine RNAiMax Transfection Reagent according to the manufacturer’s instructions. Human EPHB4 2YP cDNA was cloned into pLentiLox RSV-CMV-eGFP, which was used to generate EPHB4 2YP-encoding lentiviral particles at the University of Michigan Medical School Vector Core facility. HDLEC were infected with lentiviral particles (1:5 dilution) encoding EPHB4 2YP or control eGFP in the presence of 5 µg/ml of polybrene.

### HDLEC stimulation

HDLEC were stimulated 72 hours after siRNA transfection or viral transduction. For LSS and OSS induction, HDLEC in 6 cm diameter cell culture dishes were subject to continuous clockwise horizontal rotation (LSS) or alternating horizontal clockwise and anti-clockwise rotation for 1s each (OSS) on an orbital shaker set at 100 rpm (Major Science). The wall shear stress is estimated at 6 dynes/cm^2^. In shear stress experiments that examined the effect of AZD6244 (0.5μM), the drug was included in cultures from the start of shear stress induction. In shear stress experiments that examined the effect of GsMTx4 (5μM), all samples were treated with the drug for a total of 60 minutes. OSS was applied for all 60 minutes, the last 10 minutes of culture, or not at all. For growth factor stimulation, HDLEC in 6 well culture plates were stimulated with 100 ng/ml of VEGF-C. For Peizo1-specific stimulation, HDLEC in 6 well culture plates were stimulated with 500 nM of Yoda1.

### Western blotting

Whole-cell lysates were prepared in RIPA buffer. Protein abundance was determined by Western blotting using rabbit anti–phospho–ERK, rabbit anti-ERK, rabbit anti-β-Tubulin, mouse anti-RASA1, goat anti-EPHB4 (AF446 or AF3038), mouse anti–integrin alpha-9, rabbit anti-FOXC2, rabbit anti-actin, and mouse anti-GAPDH antibodies followed by goat anti-mouse IgG-HRP, goat anti-rabbit IgG-HRP and donkey anti-goat IgG-HRP as appropriate.

### RTqPCR

Total RNA was extracted from HDLEC using an RNeasy Micro Kit, and reverse transcribed into cDNA Synthesis using an iScript™ cDNA Synthesis Kit. *FOXC2* gene expression was analyzed by qPCR using QuantStudio™ 3 Real-Time PCR System using *GAPDH* internal controls.

## ACKNOWLEDGMENTS

We acknowledge Wanda Filipiak and Galina Gavrilina and the Transgenic Animal Model Core and Roland Hilgarth and Tonya Kopas of the Vector Core of the University of Michigan’s Biomedical Research Core Facilities for assistance with the design and production of EPHB4 2YP knockin mice and EPHB4 2YP lentiviruses respectively. Professor Taija Makinen (Uppsala University, Sweden) is thanked for providing *Ephb4* floxed mice. Drs. Sathish Srinivasan and Xin Geng (Oklahoma Medical Research Foundation), Dr. Titus Bogon (Yale University), and Dr. Taija Makinen (Uppsala University) are thanked for helpful discussions. This work was supported by NIH/NHLBI grants R01 HL146352 and R01 HL120888 to P.D.K., R01 1122578 to M.J.D.

## AUTHOR CONTRIBUTIONS

D.C. and P.D.K. designed all experiments except LV valve function tests and near-infrared imaging studies that were designed by M.J.D. and E.M.S., respectively. Experiments were conducted by D.C., D.W. and M.J.D. Data was analyzed and interpreted by D.C., P.D.K. M.J.D, and E.M.S. Funding was secured by P.D.K., M.J.D., and E.M.S. P.D.K. supervised and directed the project. The manuscript was written by P.D.K. with input from D.C., M.J.D. and E.M.S.

## DECLARATION OF INTERESTS

The authors declare no competing interests.

## INCLUSION AND DIVERSITY

We support inclusive, diverse, and equitable conduct of research.

## FIGURE LEGENDS

**Figure S1.**
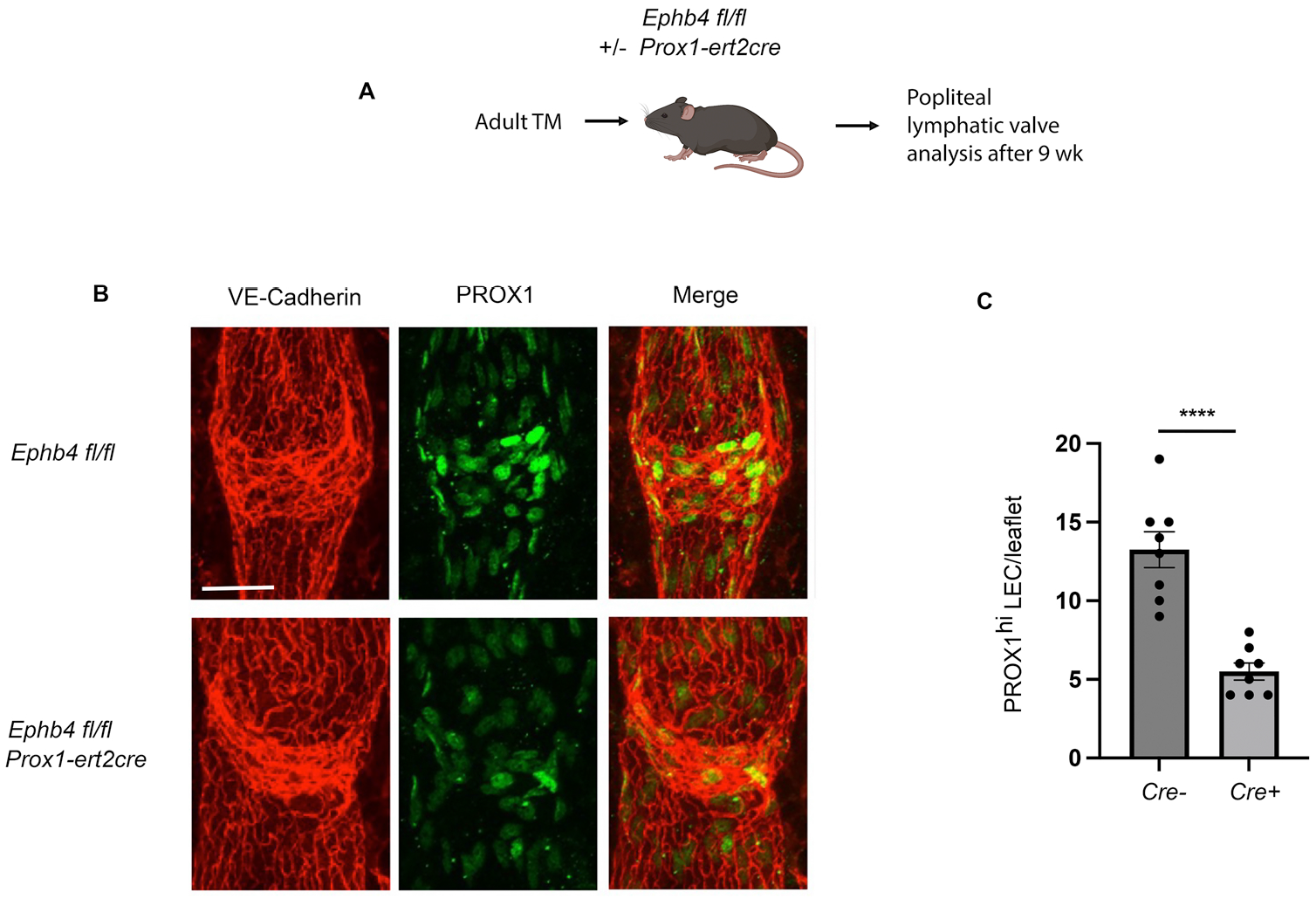
Expression of PROX1 in LV leaflets of induced LEC-specific adult EPHB4-deficient mice. (A) Schematic showing design of experiments in (B and C). (B) Representative images of collecting popliteal LV valves stained with antibodies against VE-Cadherin and PROX1. Scale bar=50 µm. (C) Mean +/− 1 SEM of PROX1^hi^ LEC per LV valve leaflet from mice of the indicated genotypes (n=8 each genotype). **** *P*<0.0001, two-tailed Student’s t-test.

**Figure S2.**
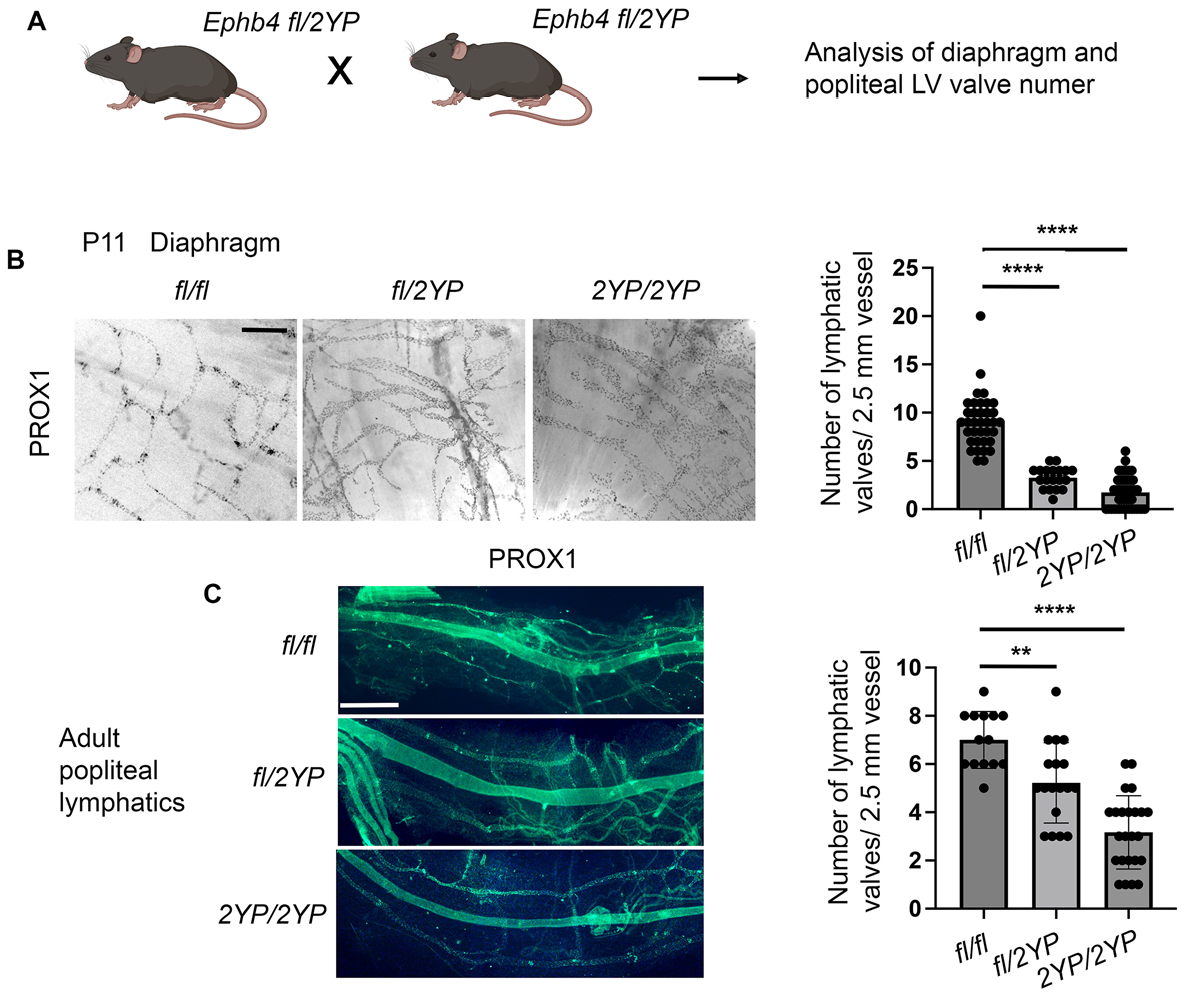
Numbers of valves in diaphragmatic and popliteal collecting LV of EPHB4 2YP mice. (A) Schematic showing design of experiments in (B and C). (B) Representative images of diaphragmatic LV from P11 pups stained with PROX1 are shown at left. Scale bar=100 µm. Graph shows mean +/− 1 SEM of the number of diaphragmatic LV valves in P11 pups of the indicated genotypes (*Ephb4 fl/fl*, n=36 determinations from 5 mice; *Ephb4 fl/2YP*, *n*=19 determinations from 3 mice; *Ephb4 2YP/2YP*, n=47 determinations from 4 mice). **** *P*<0.0001, two-tailed Mann-Whitney test. (C) Representative images of popliteal LV from adult mice stained with PROX1 are shown at left. Scale bar=100 µm. Graph shows mean +/− 1 SEM of the number of popliteal LV valves in adult mice of the indicated genotypes (*Ephb4 fl/fl*, n=14 determinations from 2 mice; *Ephb4 fl/2YP*, *n*=18 determinations from 4 mice; *Ephb4 2YP/2YP*, n=24 determinations from 4 mice). ** *P*<0.01, **** *P*<0.0001, two-tailed Student’s t-test.

**Figure S3.**
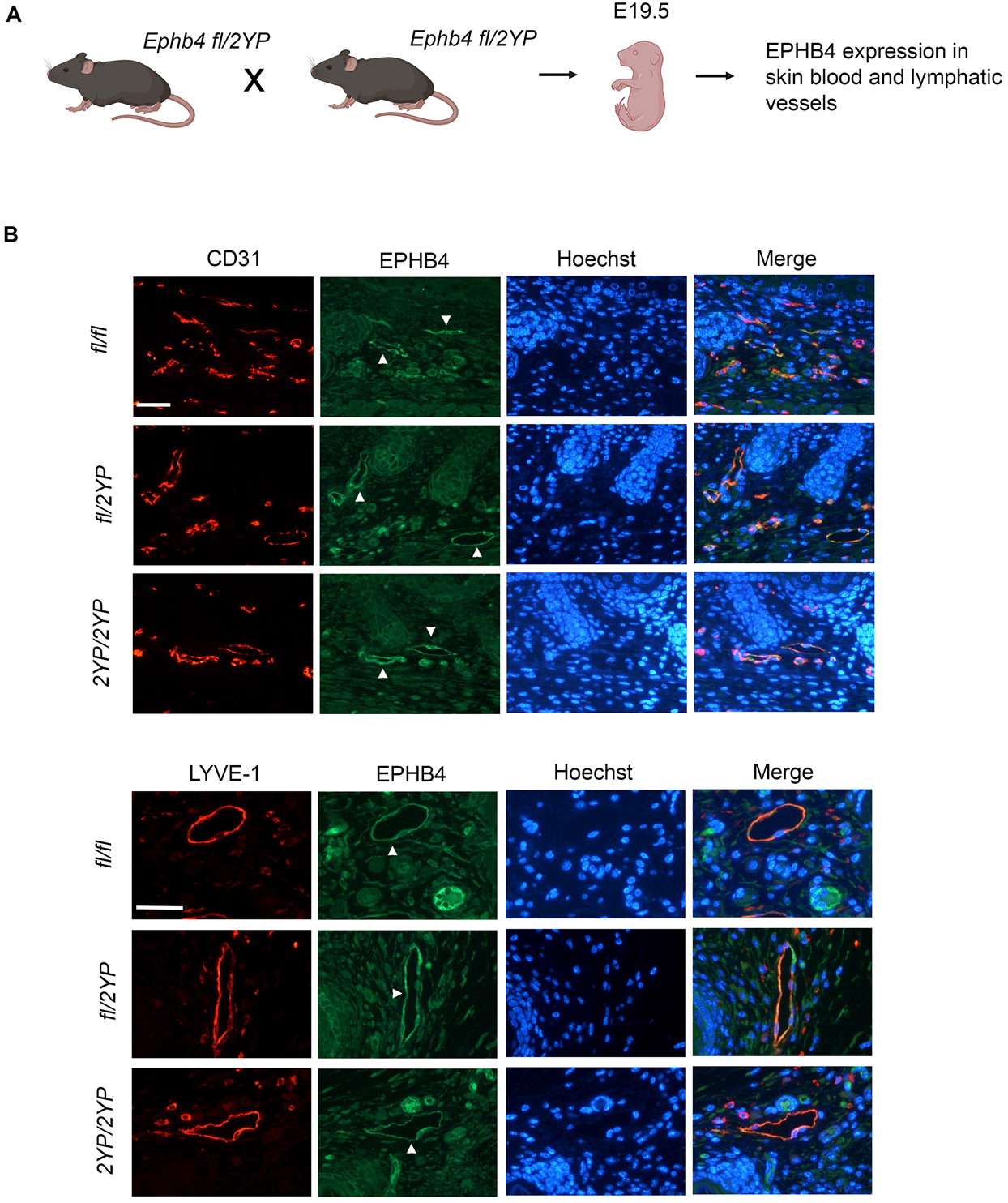
EPHB4 expression in BV and LV of EPHB4 2YP embryos. (A) Schematic showing design of experiments in (B and C). (B) Sections of skin stained with an anti-EPHB4 antibody and anti-CD31 or anti-LYVE-1 antibodies to identify BV and LV respectively (some marked with arrowheads). Scale bars=100 µm.

**Figure S4.**
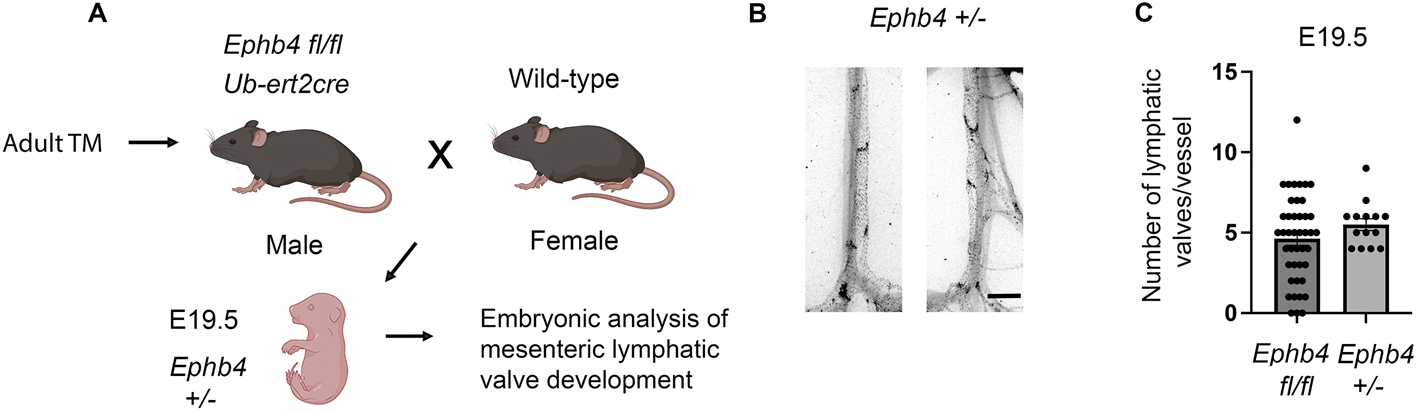
Normal LV valve development in heterozygous null EPHB4 mice. (A) Schematic showing method used to generate *Ephb4 +/−* embryos (B) At left are shown representative images of mesenteric LV from E19.5 *Ephb4 +/−* embryos stained for PROX1. Graph shows mean +/− 1 SEM of the number of valves per LV (*Ephb4 +/−*, n=14 vessels from 3 embryos; data from E19.5 *Ephb4 fl/fl* embryos without AZD6244 from Figure 3F is shown for comparison, n=43 vessels from 7 embryos). Scale bar=200 µm.

**Figure S5.**
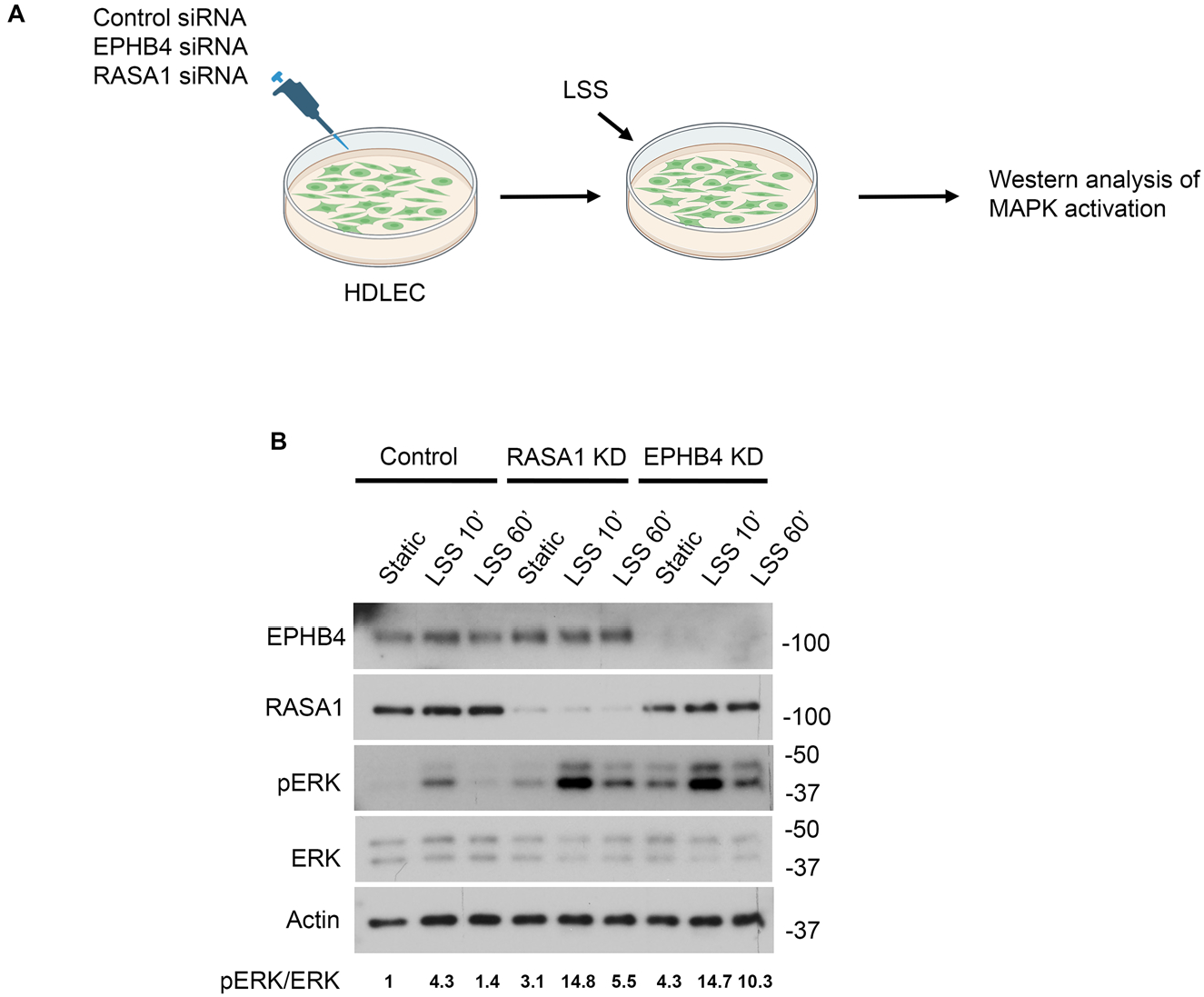
LSS-induced activation of MAPK in HDLEC. (A) Schematic showing design of experiments in (B). (B) LSS-induced MAPK activation in HDLEC treated with control, RASA1, or EPHB4 siRNA.

**Figure S6.**
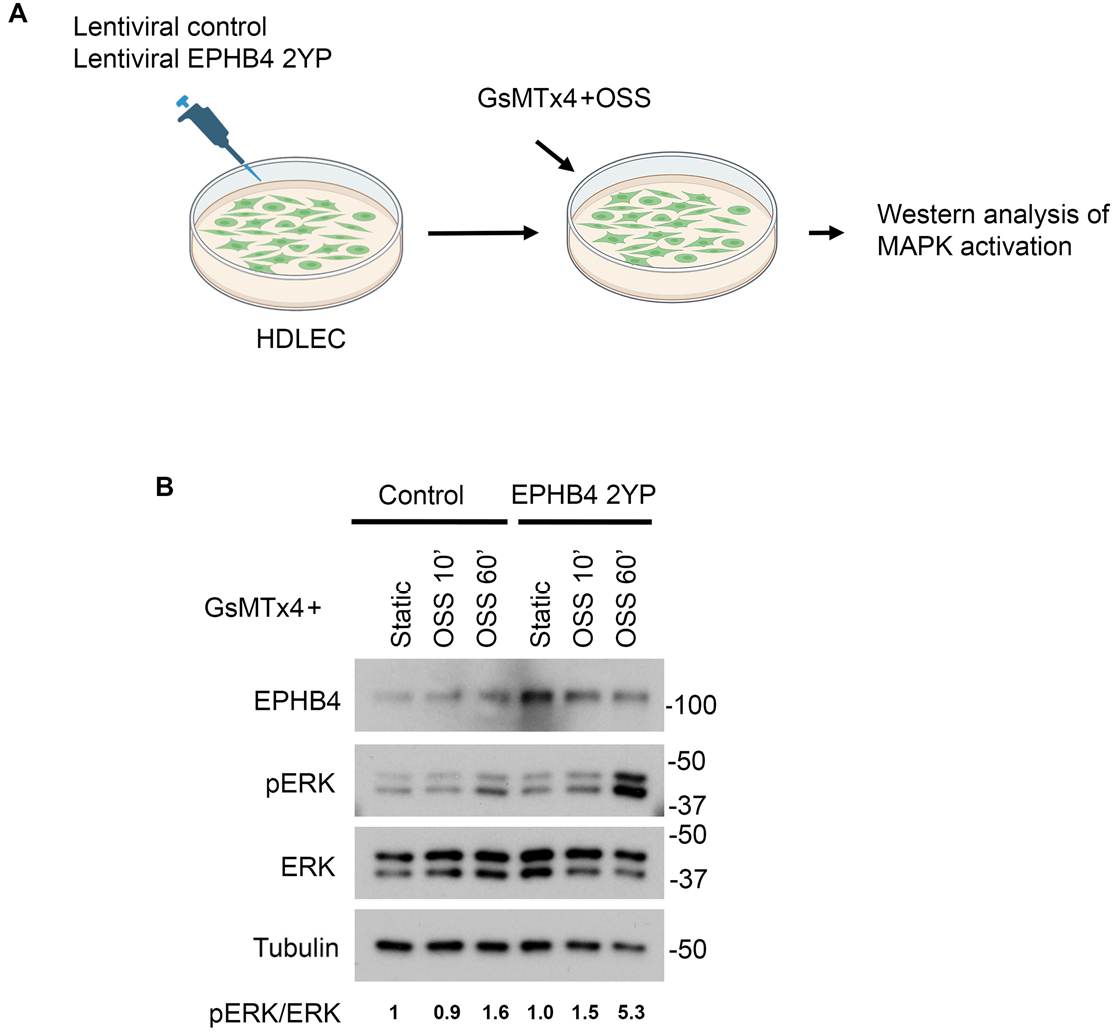
EPHB4 and RASA1 regulation of OSS-induced MAPK activation in HDLEC. (A) Schematic showing design of experiments in (B). (B) OSS+GsMTx4-induced MAPK activation in HDLEC treated with control lentivirus or lentivirus encoding EPHB4 2YP. Blots were probed with anti-tubulin antibodies to demonstrate equivalent protein loading.

## Notes

### Competing Interest Statement

The authors have declared no competing interest.

